# Hypogravity reveals a proprioception-dependent mechanism for locomotor state expression

**DOI:** 10.64898/2026.06.02.729513

**Authors:** Alessandro Santuz, Francesco Luciano, Valentina Natalucci, Adama Mbaye, Nini Ma, Dario Cazzola, Steffi Colyer, James Cowburn, Kirsten Albracht, Bjoern Braunstein, Jörn Rittweger, Nolan Herssens, Tobias Weber, David A. Green, Joriene de Nooij, Alberto E. Minetti, Gaspare Pavei, Niccolò Zampieri

## Abstract

Animals must adapt locomotion to changing environmental conditions, but how the nervous system flexibly controls movement remains unclear. Gravity provides a powerful perturbation because it alters body loading, limb dynamics, and proprioceptive feedback. Apollo astronauts often skipped on the Moon, adopting an asymmetric gait rarely used on Earth, yet the sensorimotor basis of this behaviour is unknown. Here, by studying hypogravity locomotion in humans and mice, we show that reduced gravity reveals a conserved proprioception-dependent mechanism for locomotor state expression. In humans, skipping in simulated lunar gravity is generated through flexible temporal reconfiguration of existing motor modules rather than construction of a new motor architecture. In mice, simulated lunar gravity elicited a skipping-like locomotor state, and genetic elimination of muscle proprioceptors abolished this response. Thus, hypogravity reveals how proprioceptive feedback enables flexible expression of locomotor states.

## Introduction

A central challenge in neuroscience is understanding how the nervous system controls movement in a robust and flexible way. This is especially evident during locomotion, where coordination must be maintained while the sensorimotor system adapts to changing body state and environmental conditions^1–3^. Somatosensory feedback is central to this flexibility because it continuously informs the nervous system about limb position, loading, contact, and body movement. Gravity represents a powerful perturbation of this control problem because it simultaneously alters body loading, limb dynamics, and the resulting sensory feedback^4–6^. Thus, changes in gravitational conditions offer a unique opportunity to ask how the nervous system selects among available locomotor states when the mechanical and sensory context of movement is altered.

During extravehicular activities on the lunar surface, Apollo astronauts often preferred a unilateral skipping gait over walking or running (Fig. 1A and movie S1)^7^, citing stability and ease of adaptation as reasons for their choice^8–10^. This behaviour is striking because unilateral skipping is rarely used by adults on Earth, yet it emerged spontaneously in the reduced-gravity environment of the Moon. Skipping is a hybrid gait that combines the double support of walking with the flight phase of running and involves an asymmetric use of the limbs, as the trailing and leading legs never switch roles (Fig. 1B and C)^7,11^. On Earth, skipping is metabolically more costly than running^7^. This energetic disadvantage progressively diminishes as gravity decreases, and under lunar gravity it becomes negligible^12^. Therefore, energetic viability alone does not explain why skipping becomes a preferred locomotor solution in hypogravity, an issue that is increasingly relevant as the Artemis programme aims to establish a sustained human presence on the Moon^13^.

**Fig. 1.**
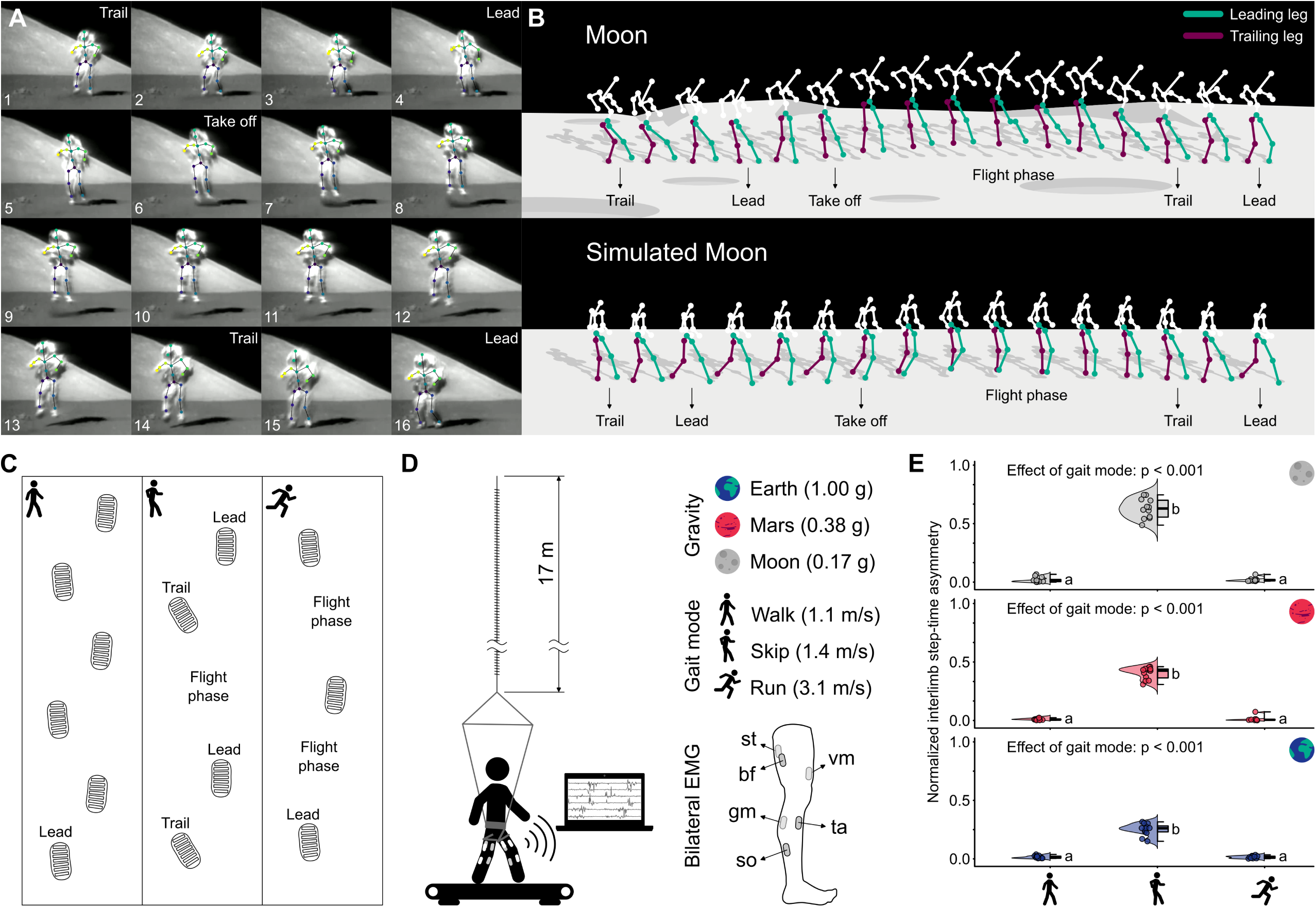
Lunar skipping is an asymmetric gait. (**A**) Two-dimensional, markerless tracking of Eugene Cernan skipping on the Moon during the Apollo 17 mission. (**B**) Sagittal-plane stick diagrams reconstructed from the Apollo archival footage (top) and from simulated hypogravity on a treadmill with body weight support (bottom). (**C**) Schematic footfall pattern of walking (left), unilateral skipping (center), and running (right). Unlike walking and running, skipping maintains the same trailing limb across cycles and combines a walking-like double stance with a running-like flight phase. (**D**) Human simulated hypogravity locomotion setup. Participants locomoted on a treadmill under Earth’s gravity, as well as under simulated Martian and lunar gravity, while supported by an elastic body-weight suspension system. Surface electromyography (EMG) was recorded bilaterally from the *vastus medialis* (vm), *semitendinosus* (st), *biceps femoris* long head (bf), *tibialis anterior* (ta), *gastrocnemius medialis* (gm), and *soleus* (so). (**E**) Interlimb step-time asymmetry across gravity conditions and gaits (*N* = 12 participants). Asymmetry was quantified as the absolute difference between lead-to-trail and trail-to-lead touchdown intervals, normalised to mean gait-cycle duration. Different letters indicate significant *post-hoc* differences.

How reduced gravity changes the neural control of locomotion remains unknown. One possibility is that locomotion in hypogravity requires a new motor control architecture. An alternative is that reduced gravity reveals a latent locomotor state by retuning pre-existing control elements. Muscle synergies describe the modular organisation of movement and provide a way to distinguish between these two possibilities^14^. A new control architecture should alter the relative contribution of muscles to each synergy. Reuse of existing synergies should instead preserve the modular structure while changing the timing of recruitment^15–20^. Another unresolved question is whether the effect of hypogravity depends on sensory feedback. Proprioception, the sense of body position and movement, monitors muscle length and tension and is therefore ideally positioned to signal the changes in limb loading produced by reduced gravity^21^. If the emergence of skipping on the Moon reflects a sensory-dependent adaptation, then disrupting proprioceptive feedback should impair the expression of the asymmetric locomotor state induced by hypogravity. Answering these questions requires an experimental model in which locomotion can be tested under reduced gravity and the sensorimotor system can be causally manipulated.

We therefore combined simulated hypogravity experiments in humans with a newly developed mouse model of reduced-gravity locomotion. In humans, muscle-synergy analysis allowed us to test whether hypogravity changes the structure or timing of locomotor control. In mice, locomotion under simulated lunar gravity allowed us to determine whether reduced gravity induces an adaptive locomotor state and to test whether proprioceptive feedback is required for its expression. Together, this cross-species approach reveals a proprioception-dependent mechanism governing expression of locomotor states.

## Results

### Simulated lunar gravity reproduces the asymmetric structure of skipping

We modelled human reduced-gravity locomotion in the laboratory using an elastic body-weight support system mounted over a treadmill. Participants were asked to walk, skip, and run under Earth gravity and simulated Martian and lunar gravity while bilateral electromyographic (EMG) activity was recorded from lower-limb muscles (Fig. 1D and movie S2). Stick-figure reconstruction of skipping obtained during simulated lunar gravity recapitulated the kinematic pattern observed in the Apollo footage (Fig. 1B), indicating that the experimental paradigm captured the essential spatiotemporal organisation of lunar skipping. Quantitative analysis confirmed that skipping was the gait characterised by the strongest asymmetry across all gravity conditions (Fig. 1E). Interlimb step-time asymmetry, computed as the normalised absolute difference between lead-to-trail and trail-to-lead touchdown intervals, was higher for skipping than for either walking or running, and this feature was conserved under simulated Martian and lunar gravity.

These results establish that simulated hypogravity reproduces the interlimb asymmetry characteristic of lunar skipping.

### Hypogravity reconfigures the timing of conserved locomotor modules

To determine whether locomotion in hypogravity requires a new control architecture or the retuning of an existing one, we extracted muscle synergies^14^ from lower-limb EMG activity (Fig. S1) recorded during walking, skipping, and running under Earth gravity and simulated Martian and lunar gravity. Muscle synergies are composed of time-independent muscle weights (**W**) and time-dependent activation patterns (**P**) describing, respectively, the relative contribution of different muscles to each synergy and the time course of synergy activation (Fig. 2A). The number of synergies (Fig. 2B) and quality of EMG reconstruction (Fig. 2C) were unaffected by gait or gravity, indicating that neither parameter alters the dimensionality of motor output. Four muscle synergies functionally described different phases of the gait cycle in walking, skipping, and running (Fig. 2D and Table S1). Activation patterns (**P**) showed pronounced temporal modulation in response to hypogravity (Fig. 2D), such as shifts (Fig. 2E) and/or changes in the duration of muscle synergy activity (Fig. S2). This is consistent with the idea that adaptation to perturbation is primarily achieved by adjusting the timing of a conserved modular architecture^15–20^.

**Fig. 2.**
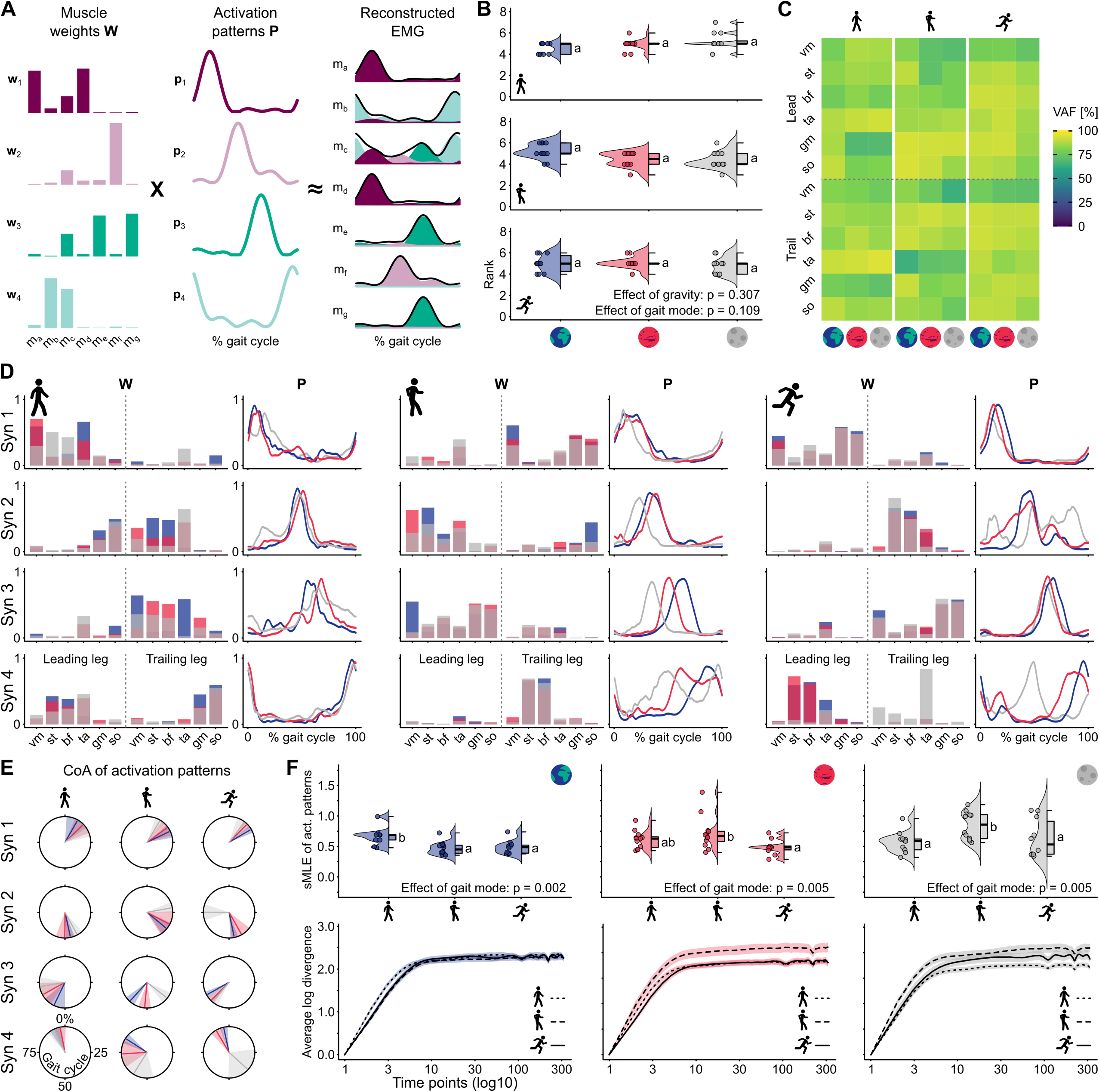
Hypogravity induces the modulation of muscle synergy activation patterns. (**A**) Mathematical intuition of non-negative matrix factorisation of EMG activity to extract muscle synergies. Each synergy is defined by a set of time-independent muscle weights **W**, representing the relative contribution of each muscle within a synergy, and a set of time-dependent activation patterns **P**, representing the time course of synergistic activation. (**B**) Minimum number of synergies (factorisation rank) required to reconstruct locomotor EMG patterns across gaits and gravity conditions. (**C**) Reconstruction quality of the synergy model, expressed as variability accounted for (VAF), across recorded muscles, gaits, and gravity conditions. (**D**) Muscle weights **W** and activation patterns **P** of muscle synergies extracted from walking, skipping, and running under Earth gravity, and under simulated Martian and lunar gravity. (**E**) Center of activity (CoA) of synergy activation patterns across gaits and gravities. (**F**) Local dynamic stability of activation patterns, quantified by the short-term maximum Lyapunov exponent (sMLE), across gaits and gravities (*N* = 12 participants). Different letters indicate significant *post-hoc* differences.

Because hypogravity affected the activation patterns of a conserved number of muscle synergies, we next asked whether it also altered their dynamical properties. Previous work showed that perturbations make muscle activation patterns less unstable during locomotion^22^, suggesting that challenging conditions constrain the range of activation dynamics. By contrast, less constrained conditions may permit a wider range of activation solutions. We therefore calculated the short-term maximum Lyapunov exponent of activation patterns to quantify their dynamic stability^23^. The results show that, on Earth, the gait with the largest instability of activation patterns was walking (Fig. 2F). Yet, under simulated lunar gravity, skipping presented the highest instability (Fig. 2F). Thus, hypogravity reorganised activation pattern dynamics, shifting the greatest instability from walking on Earth to skipping under lunar gravity, consistent with skipping entering a more flexible activation-dynamics regime under reduced gravity.

Altogether, these findings indicate that hypogravity primarily alters locomotor control by changing the timing of a conserved set of muscle synergies, rather than by changing the modular architecture of the synergies.

### Skipping shares a running-like modular architecture

To determine whether skipping arises through reuse of pre-existing locomotor modules, we compared muscle synergies extracted from skipping EMG with those extracted from walking and running. Skipping and running, unlike walking, are bouncing gaits^7,12^ and, although kinematically distinct, their temporal activation patterns under Earth gravity were qualitatively similar (Fig. 3A). By contrast, the corresponding muscle weights, ordered according to the timing of their activation patterns, differed substantially (Fig. 3B). However, reordering the skipping muscle weights from their original sequence 1-2-3-4 to a new sequence of recruitment (3-4-1-2) revealed a marked overlap with the muscle weights of running (Fig. 3C). These observations suggest that the main difference between skipping and running does not lie in the structure of activation patterns, but in the recruitment order of muscle weights.

**Fig. 3.**
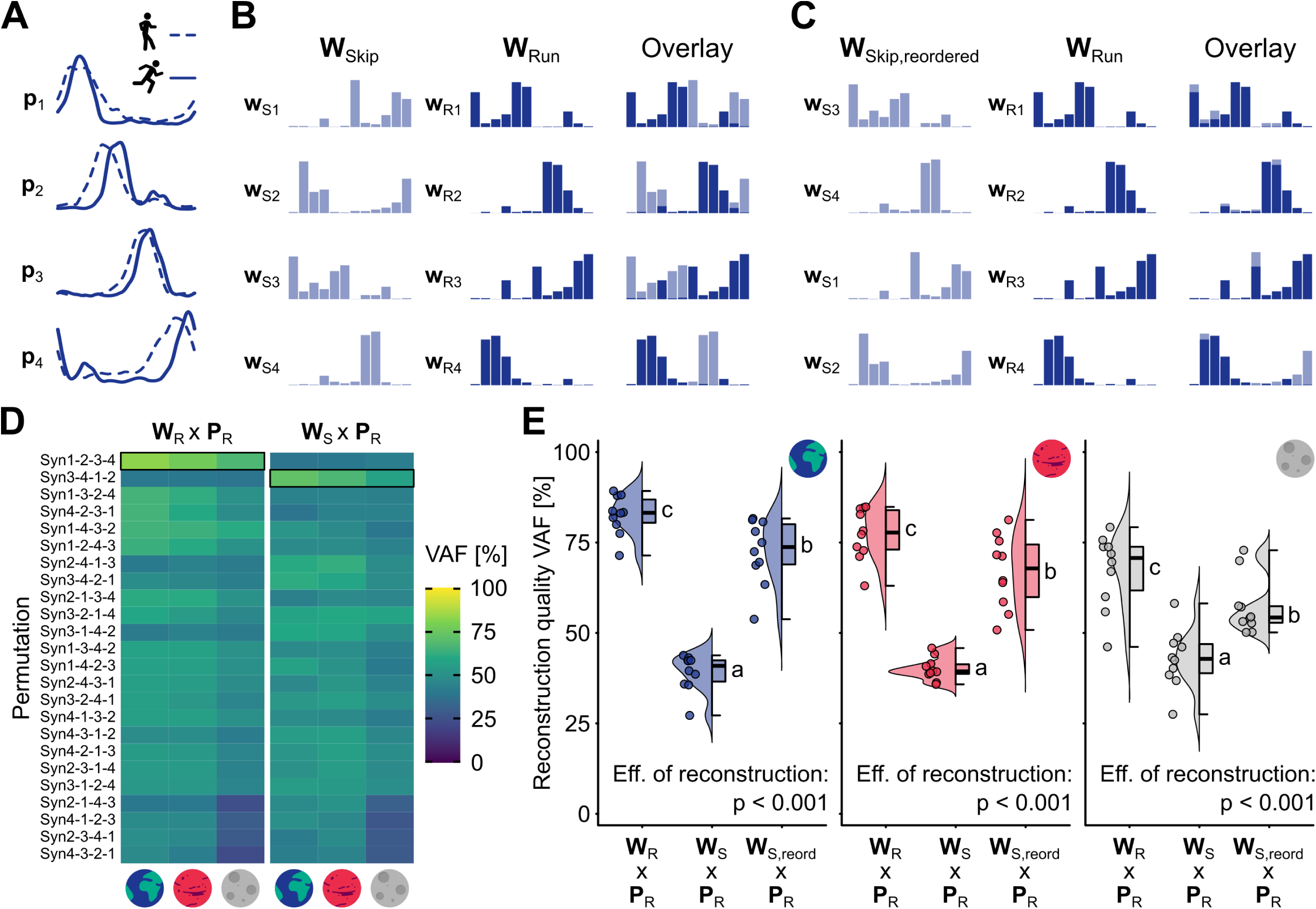
Skipping emerges from a running-like modular architecture. (**A**) Average synergy activation patterns **P** for running and skipping under Earth gravity. (**B**) Synergy muscle weights **W** for running and skipping under Earth gravity. (**C**) Reordered skipping muscle weights overlaid on running weights. (**D**) Cross-reconstruction analysis of running EMG using running activation patterns combined with running weights or skipping weights. Reconstruction quality was quantified for all possible weight-order permutations. (**E**) Reconstruction quality of running across gravity conditions using i) running weights and activations, ii) skipping weights and running activations, or iii) reordered skipping weights and running activations. Different letters indicate significant *post-hoc* differences (*N* = 12 participants).

To test this idea systematically, we performed a cross-reconstruction analysis in which running EMG activity was reconstructed using all possible permutations of running activation patterns and muscle weights from either running or skipping (**W** x **P**; Fig. 3D). As expected, running EMG was best reconstructed by using running muscle weights ordered according to original timing of running activation patterns (1-2-3-4), but reconstruction quality decreased when the temporal sequence of running muscle weights recruitment was changed (Fig. 3D, left). Conversely, reconstruction of running EMG using skipping muscle weights ordered using the original sequence of skipping activation patterns was poor, consistent with a mismatch in the spatial structure of running and skipping (Fig. 3D, right). However, reordering the skipping muscle weights specifically into the 3-4-1-2 sequence yielded high reconstruction quality across gravity levels (Fig. 3E). This effect was not observed when the same analysis was applied to walking and skipping (Fig. S3).

These results indicate that skipping is generated by reconfiguring a running-like architecture rather than by using a new set of locomotor modules.

### Simulated lunar gravity induces a skipping-like locomotor state in mice

To determine whether the emergence of skipping is a conserved adaptation to hypogravity and identify the underlying mechanisms, we established a mouse model of reduced-gravity locomotion. We adapted the human experimental paradigm to mice by studying animals wearing a harness connected to an elastic body-weight support system over a treadmill (Fig. 4A). Control experiments showed that although the harness reduced spontaneous exploration in an open-field assay, it did not affect their preferred speed of locomotion (Fig. S4A). The attachment point of the elastic on the harness was aligned with the calculated centre of mass (50.2% of body length; Fig. S4B), minimizing rotational moments during body-weight support. High-speed video and markerless motion tracking (Fig. 4A) were used to quantify 134 kinematic parameters (Table S2) during locomotion under Earth and simulated lunar gravity^24^. Unlike in the human experiments, mice spontaneously selected their gait.

**Fig. 4.**
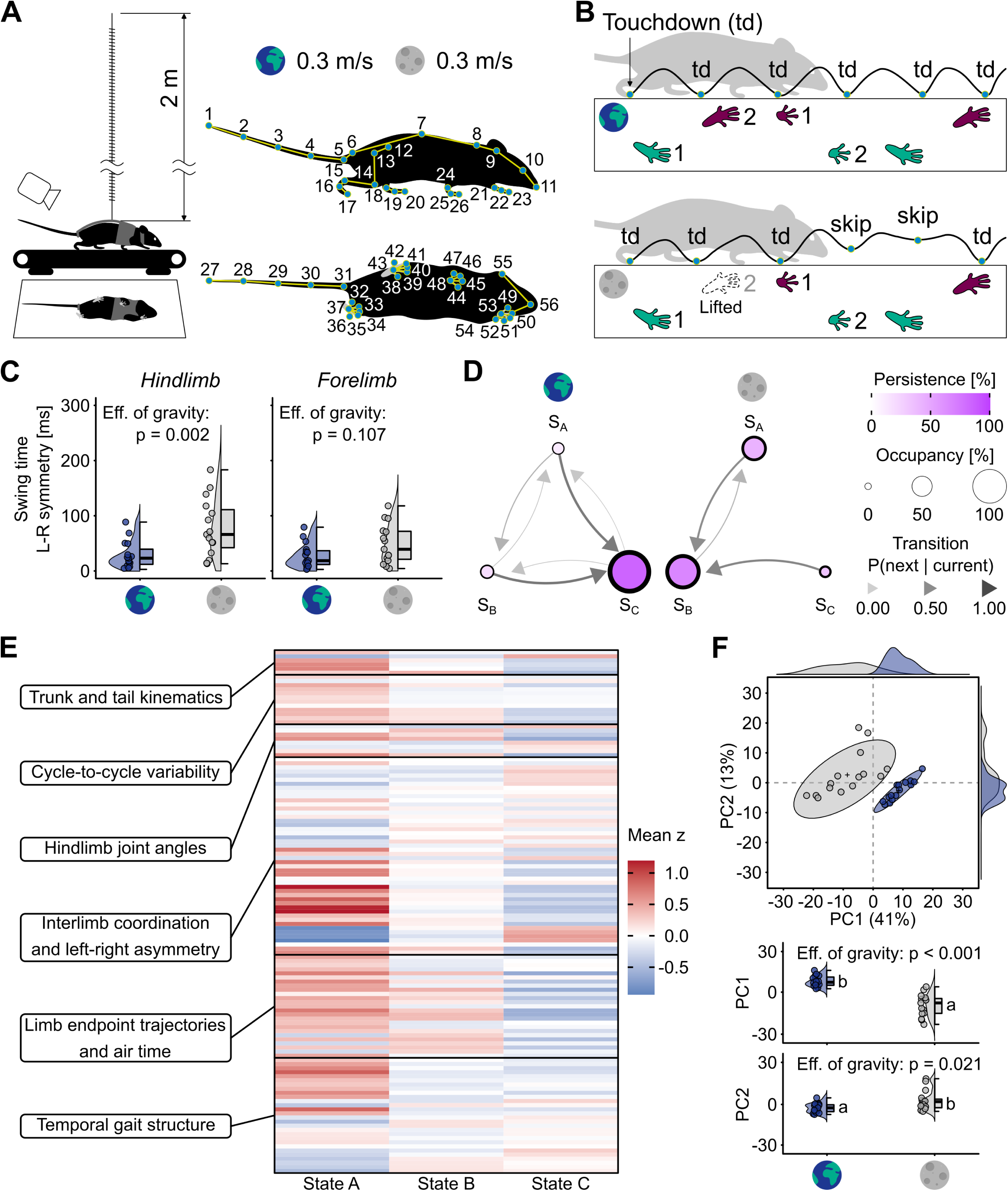
Hypogravity induces skipping-like asymmetric gait in mice. (**A**) Mouse simulated hypogravity locomotion setup. Mice walked on a treadmill at 0.3 m/s under Earth gravity or simulated lunar gravity while supported by an elastic body-weight suspension system. Limb kinematics were quantified from high-speed video using markerless tracking. (**B**) Representative limb trajectories under Earth gravity and simulated lunar gravity. Under Earth gravity, mice typically used a symmetric trot (top). Under simulated lunar gravity, mice frequently prolonged swing of one hindlimb across one or more gait cycles, producing a persistent hindlimb asymmetry (bottom). (**C**) Left–right airtime asymmetry during Earth and simulated lunar gravity locomotion for hindlimbs and forelimbs (*N* = 15 mice). (**D**) Locomotor state analysis during Earth and simulated lunar gravity locomotion. State-transition graphs summarise the three locomotor states (S_A_, S_B_, and S_C_) identified by unsupervised clustering. Node size indicates state occupancy, node color indicates state persistence (probability of remaining in the same state across consecutive strides), and arrow thickness indicates transition probability between states. (**E**) Heat map of z-scored kinematic variables defining the locomotor states. Variables were grouped according to trunk and tail kinematics, cycle-to-cycle variability, hindlimb joint angles, interlimb coordination and left–right asymmetry, limb endpoint trajectories and airtime, and temporal gait structure. Mean normalised values for each variable are shown for S_A_, S_B_, and S_C_. (**F**) Principal component analysis of kinematic variables during Earth and simulated lunar locomotion (*N* = 15 mice). Different letters indicate significant *post-hoc* differences.

Under Earth gravity, mice typically locomoted with a symmetrical trot (Fig. 4B and movie S3)^25^. In contrast, under simulated lunar gravity, the animals frequently adopted asymmetric limb movements characterised by prolonged swing duration of one limb relative to the other, such that one paw remained lifted for one or more gait cycles (Fig. 4B and movie S3). Although quadrupedal in expression, this behaviour was analogous to unilateral skipping in humans because it introduced pronounced left-right asymmetry in hindlimb timing. To quantify this effect, we calculated left-right airtime symmetry for both forelimbs and hindlimbs. Simulated lunar gravity increased asymmetry in the hindlimbs, whereas the forelimbs were not significantly affected (Fig. 4C).

To determine whether these changes reflected a broader reorganisation of locomotion, we analysed recurrent patterns in gait kinematics using hidden Markov modelling. This analysis identified three recurrent locomotor states whose persistence, occupancy, and transition probability differed markedly between Earth and simulated lunar gravity, indicating that hypogravity reorganises the expression of distinct locomotor states (Fig. 4D). States were statistically separated by several locomotion features, including trunk and tail kinematics, hindlimb joint angles, temporal gait structure, limb trajectories, and interlimb coordination (Fig. 4E, S4C). This demonstrates that hypogravity reorganises locomotion across multiple levels of motor control. Consistent with this result, principal component analysis of the full kinematic dataset revealed significant separation between Earth and simulated lunar gravity, supporting the conclusion that reduced gravity induces a distinct locomotor state in mice (Fig. 4F).

These findings demonstrate that key features of the locomotor response to hypogravity are conserved across mammals and provide a tractable model system for identifying the sensory mechanisms that govern its expression.

### Progressive proprioceptor loss causes graded locomotor impairment

The emergence of a skipping-like locomotor state under lunar gravity provides a powerful experimental model for testing the sensory mechanisms that enable this adaptation. Proprioception is ideally positioned to detect changes in gravitational load because muscle spindles and Golgi tendon organs continuously monitor muscle length and tension, thereby signalling how limb mechanics are altered when gravity changes.

To test whether proprioceptive feedback is required for the emergence of this locomotor adaptation, we selectively ablated proprioceptive sensory neurons. Previous studies have targeted proprioceptive afferents by taking advantage of Parvalbumin (PV) expression in dorsal root ganglia^26,27^. However, because PV is also expressed by a subset of low-threshold cutaneous mechanoreceptors, this approach cannot isolate the contribution of proprioceptive from cutaneous sensory feedback^28,29^. To overcome this limitation, we used an intersectional genetic strategy exploiting the co-expression of PV and Runx3 (Rx3) to selectively target muscle spindle and Golgi tendon organ afferents^30^ for acute ablation using the diphtheria toxin (DT) system^31^. We generated mice in which *PV::Cre* and *Rx3::FlpO* drive the expression of the human DT receptor and a red fluorescent protein (Fig. 5A, S5). Following DT administration in adult mice, histological analysis of dorsal root ganglia revealed progressive loss of tdTomato-labelled proprioceptors beginning 24 hours after injection and reaching maximal levels by 72 hours, with no further significant reduction up to one week (Fig. 5B).

**Fig. 5.**
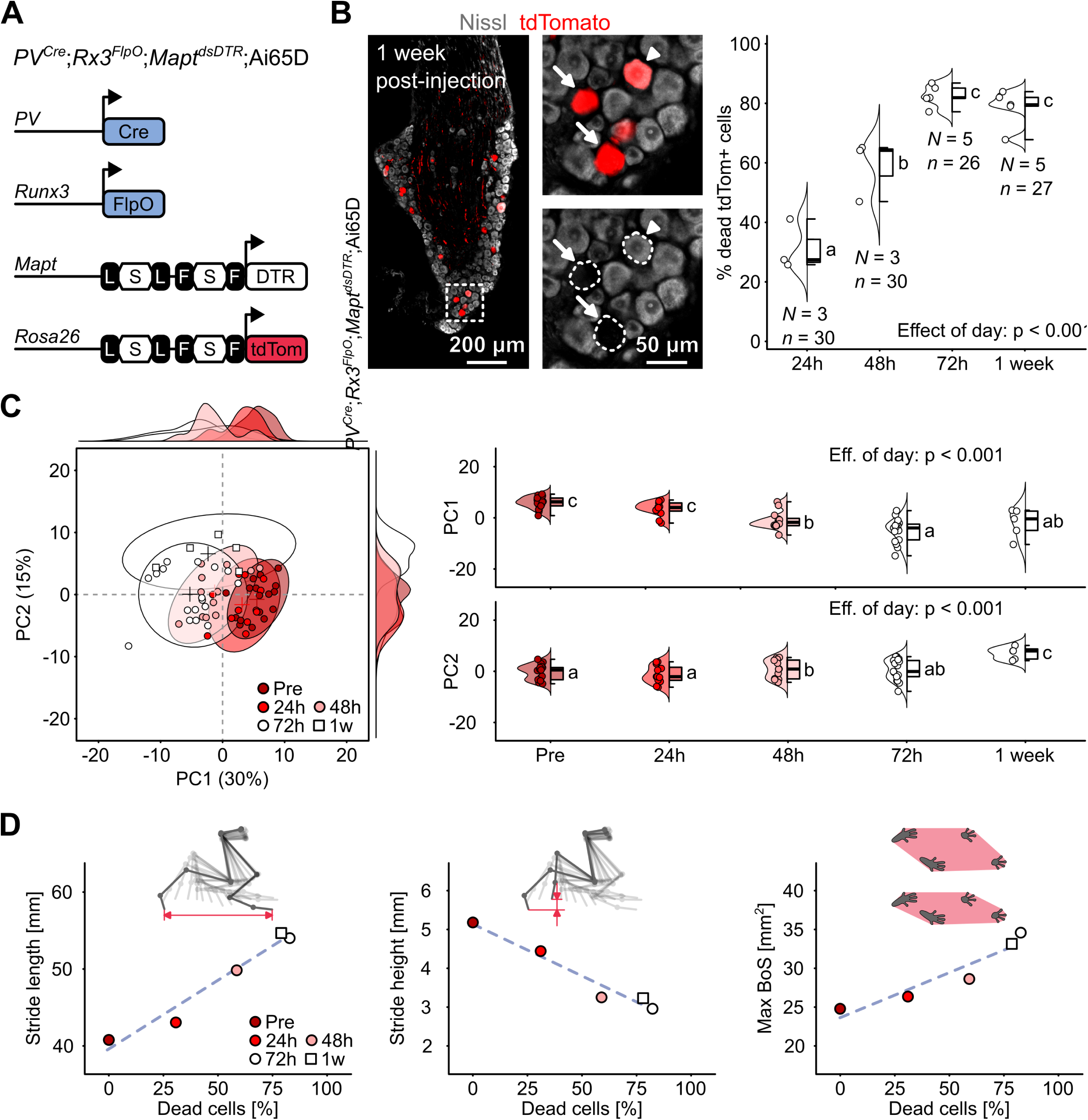
Progressive elimination of proprioceptive afferents causes graded locomotor impairments. (**A**) Intersectional genetic strategy combining Parvalbumin and Runx3 expression to selectively express diphtheria toxin receptor (DTR) and tdTomato in proprioceptive sensory neurons. (**B**) Representative image of a lumbar dorsal root ganglion one week after diphtheria toxin administration showing loss of proprioceptive afferents (left, arrows indicate ablated neurons, tdTomato^+^; Nissl^-^; arrowhead shows a spared neuron, tdTomato^+^; Nissl^+^) and time course of neuronal ablation (right). (**C**) Principal component analysis of kinematic variables before and after diphtheria toxin injection. (**D**) Relationship between proprioceptor loss and kinematic changes, using as representative examples three key parameters describing locomotion: stride length (left), stride height (center), and base of support (right).

We next asked how gradual proprioceptor ablation affected locomotion by performing a longitudinal assessment of motor performance on a treadmill. Principal component analysis of kinematic parameters (Table S2) revealed progressive separation between experimental time points beginning 48 hours after DT injection, with the largest changes occurring after 72 hours (Fig. 5C), indicating that locomotor impairment closely followed the time course of proprioceptor ablation. Across animals, changes in representative locomotor parameters scaled linearly with the extent of proprioceptor loss before reaching a plateau after 72 hours, revealing a graded dependence of locomotor performance on proprioceptive input (Fig. 5D). Locomotion remained coordinated up to 48 hours after DT administration (Movie S4), whereas severe impairment became evident only after 72 hours (Movie S5). Thus, gradual loss of proprioceptors occurring in the first 48 hours after DT injection created an experimental window in which substantial proprioceptor ablation had occurred while overall locomotor function remained largely preserved, allowing us to specifically test whether proprioceptive feedback is required for hypogravity-induced adaptation before the onset of severe locomotor impairment.

### Proprioceptive feedback is required for locomotor-state adaptation to hypogravity

We next asked whether progressive loss of proprioceptive feedback alters the locomotor state evoked by hypogravity. To address this question, we examined treadmill locomotion under simulated lunar gravity before and during the first 48 hours after DT administration: the experimental window during which proprioceptor loss occurred without major disruption of locomotion.

To determine whether the organisation of the hypogravity-induced locomotor state was altered, we analysed its dynamics. Before DT administration, mice consistently occupied a distinct locomotor state associated with simulated lunar gravity (Fig. 4D, 6A). This state remained largely preserved 24 hours after DT administration but was reorganised after 48 hours, when state occupancy shifted and transition dynamics changed, indicating a disruption of the hypogravity-specific locomotor state (Fig. 6A, S6). Consistent with this reorganisation, principal component analysis of kinematic parameters revealed a separation from the pre-ablation state only at 48 hours after DT administration. Notably, this separation occurred exclusively along the second principal component, which accounted for 15% of the total variance (Fig. 6B). By contrast, locomotor patterns recorded 24 hours after ablation clustered with pre-treatment values, indicating that the hypogravity-induced locomotor state remained largely intact during the early phase of proprioceptor loss.

**Fig. 6.**
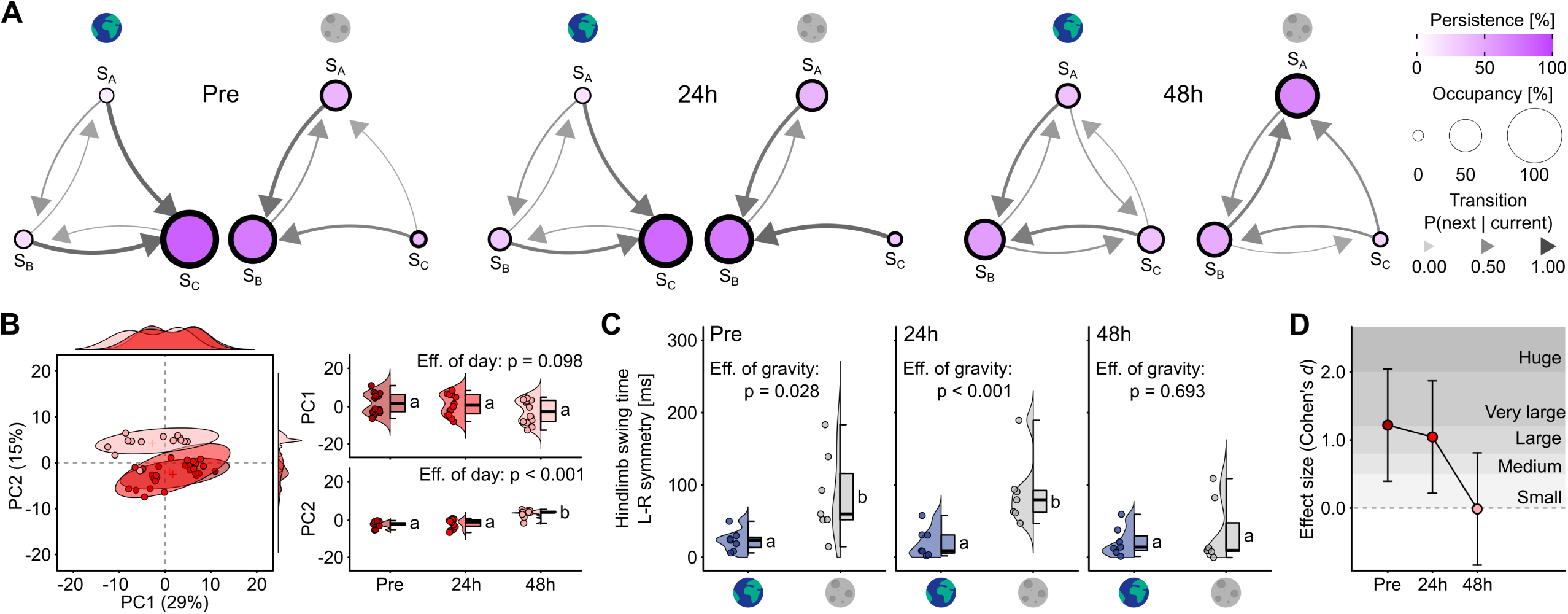
Proprioceptive feedback is required for locomotor adaptation to hypogravity. (**A**) Locomotor state analysis under Earth and simulated lunar gravity before, 24 hours, and 48 hours after proprioceptor ablation. Network graphs summarise the three locomotor states (S_A_, S_B_, and S_C_) identified by unsupervised state analysis. Node size indicates state occupancy, node color indicates state persistence (probability of remaining in the same state across consecutive strides), and arrow thickness indicates transition probability between states. (**B**) Principal component analysis of kinematic variables before and after diphtheria toxin injection. (**C**) Hindlimb left-right swing-time asymmetry under Earth and simulated lunar gravity before (left), 24 hours (center) and 48 hours (right) after proprioceptor ablation (*N* = 7 mice). (**D**) Effect size of simulated lunar gravity on hindlimb asymmetry over time after diphtheria toxin injection (*N* = 14 mice). Different letters indicate significant *post-hoc* differences.

We therefore asked whether this change in the hypogravity-induced locomotor state was accompanied by loss of the characteristic hindlimb asymmetry. Before DT administration, and still 24 hours after injection, simulated lunar gravity induced asymmetric hindlimb movement. This response was abolished after 48 hours and the animals adopted a symmetric gait reminiscent of that seen on Earth (Fig. 6C and movie S6). Consistent with this observation, the effect size of simulated lunar gravity on hindlimb asymmetry progressively declined from large values before and 24 hours after DT administration to negligible values at 48 hours (Fig. 6D).

Together, these findings demonstrate that proprioceptive feedback is required for the expression of the hypogravity-induced locomotor state.

## Discussion

Changes in gravity perturb the mechanical and sensory context in which locomotion is controlled: limb loading is reduced, body dynamics are reshaped, and the proprioceptive consequences of each step are altered^1^. For these reasons, the preference of Apollo astronauts for skipping on the Moon offers a unique perspective on a broad motor control question: how does the nervous system flexibly adapt locomotion when the environment constrains the available control solutions? Our findings suggest that this adaptation does not require a new motor control architecture. In humans, simulated Martian and lunar gravity preserved the number and composition of muscle synergies, while selectively changing their temporal recruitment. Thus, reduced gravity does not reshape the composition of muscles engaged in different synergies, but rather changes the temporal phase of their recruitment within the gait cycle. From this perspective, skipping is a latent locomotor state that becomes accessible when reduction in gravitational load on the Moon perturbs the sensorimotor system.

Previous studies have defined skipping as a hybrid gait that combines the double support of walking with the flight phase of running^7,11^. Our analyses extend this concept from biomechanics to motor control. Running muscle activity can be accurately reconstructed when running activation patterns are combined with an appropriate assignment of skipping muscle weights. Thus, skipping appears to arise through the flexible reconfiguration of an existing locomotor architecture rather than the recruitment of a new set of muscle synergies. Comparative observations are consistent with this view. Jerboas frequently use hopping and skipping^32,33^; crows and magpies can produce running-like mechanics through altered interlimb phasing^32,34^. Within primates, lemurs can use unilateral skipping or hopping despite lacking a human-like running gait^35^, suggesting that asymmetric bouncing patterns are part of a broader locomotor repertoire across vertebrates. In humans, children spontaneously express skipping during development^7^, indicating that this gait remains available even though it is rarely used in adulthood.

The mouse experiments in simulated lunar gravity provide a model in which the sensory mechanisms underlying this flexible adaptation can be tested. While the changes in locomotor behaviour of mice cannot be considered kinematically homologous to human skipping, simulated lunar gravity induced the key organisational feature observed in Apollo astronauts: persistent left-right asymmetry in limb timing. Under Earth’s gravity, mice locomoted with a symmetric trot. However, under simulated lunar gravity, they adopted a hindlimb-asymmetric state, in which one hind paw remained in swing for one or more gait cycles. This behaviour emerged during otherwise coordinated treadmill locomotion and affected primarily the hindlimbs, indicating that hypogravity did not disrupt locomotor control, but instead revealed an asymmetric state that parallels the defining kinematic feature of human skipping on the Moon.

We used this framework to test whether proprioception is necessary for flexible locomotor adaptation to reduced gravity by selectively eliminating muscle spindles and Golgi tendon organs afferents. Previous studies have established that proprioception is necessary for robust control of locomotor output in the face of internal or external perturbations^1,2,27,36^. However, interpretation of these experiments has been complicated because the commonly used genetic strategies rely solely on Parvalbumin expression^26,27^, which labels not only muscle proprioceptors but also approximately 10-15% of low-threshold cutaneous mechanoreceptors^28,29^. By using an intersectional strategy based on the combined expression of Parvalbumin and Runx3, we selectively manipulated proprioceptive afferents while sparing the vast majority of cutaneous mechanoreceptors. Our findings therefore provide the first direct evidence that muscle proprioceptive feedback itself is specifically required for locomotor adaptation to an external perturbation such as hypogravity.

The progressive time course of proprioceptor ablation allowed us to examine changes in hypogravity-induced locomotor state during a period in which overall locomotor performance remained largely preserved despite the elimination of approximately 60% of proprioceptive neurons. Under these conditions, the asymmetric state induced by simulated lunar gravity disappeared. Because this loss occurred before severe locomotor impairment, it cannot be explained by ataxia or global motor deterioration alone. These findings further indicate that proprioception contributes not only to the robustness of locomotor control but also to its flexibility. Notably, these functions could be dissociated: overall locomotion remained largely preserved despite substantial proprioceptor loss, whereas the adaptive locomotor response to hypogravity was abolished. These observations are consistent with a broader view in which sensory feedback is not merely corrective, but can actively shape motor control by regulating which locomotor states can be expressed.

Why does hypogravity reveal a latent locomotor state? Reduced gravity changes the physical constraints under which movement is controlled. On land, locomotion requires animals to coordinate propulsion while maintaining body support, making limb loading, stance-phase control, and balance central determinants of locomotor control^1,3,37^. Lunar gravity partially relaxes these constraints while preserving the need to coordinate propulsion, balance, and ground contact. Previous metabolic measurements showed that skipping is more costly than running on Earth, but that this energetic disadvantage becomes negligible under simulated lunar gravity. Thus, hypogravity may favour skipping not by making it metabolically cheaper than running, but by reducing the energetic penalty that normally suppresses it and allowing other factors to influence gait selection^12^. By combining an aerial phase with double support, skipping may extend the window over which body motion can be corrected before initiating the flight phase^12^. In this view, reduced gravity shifts locomotion into a regime in which a latent asymmetric bouncing solution becomes viable, while proprioceptive feedback provides the limb load- and position-dependent information required for its expression.

More broadly, our findings identify gravity as a powerful experimental paradigm for uncovering fundamental principles of sensorimotor control. Hypogravity exposes control strategies that remain largely inaccessible under terrestrial conditions. Rather than constructing a new motor programme, the nervous system flexibly accesses an existing locomotor state through proprioception-dependent reconfiguration of a conserved modular architecture. This framework links the modular organisation of motor output to sensory-dependent selection of locomotor states and provides a mechanistic explanation for one of the most iconic observations from lunar exploration.

## Methods

### Animal experiments

Human experiments were reviewed and approved by the Ethics Committees of the University of Bath (ID: EP 18/19 018) and the University of Milan (ID: 12/22). In accordance with the Declaration of Helsinki, all participants provided written informed consent prior to the study after receiving all necessary explanations. Mouse experiments were performed in accordance with European Research Council directives and were approved by the Regional Office for Health and Social Affairs Berlin (LAGeSo) under experimental license number G 0091/22. In all our experiments, both male and female mice were used. Animals were fed *ad libitum* and maintained under standard conditions on a 12-hour light/dark cycle.

### Apollo 17 extravehicular activity (EVA) tracking and 3D pose reconstruction

To analyse the Eugene Cernan’s gait, we used the video clip from the Apollo Lunar Surface Journal of Geology Station 5 at Camelot Crater during EVA 2. This clip corresponds to the segment from mission time 146:50:03 to 146:50:53. During this time, Eugene Cernan spontaneously switches to a skipping gait while returning to the Lunar Roving Vehicle. We first used DeepLabCut v2.3.9^24^ for markerless body part tracking and labelled 17 body landmarks on 40 frames taken from the original video, assigning 95% of the images to the training set without cropping. Specifically, we labelled the pelvis, right hip, right knee, right ankle, left hip, left knee, left ankle, spine, thorax, nose, head, left shoulder, left elbow, left wrist, right shoulder, right elbow, and right wrist. We trained the model using a ResNet-50-based neural network^38,39^ with default parameters for 500,000 training iterations. We validated with one shuffle and found a test error of 6.36 pixels and a train error of 1.98 pixels. We then analysed the video with this model to get 2D landmark coordinates. Because the original footage from 1972 was monocular and not purely sagittal, the resulting 2D trajectories could not be directly interpreted as sagittal-plane kinematics. Therefore, we fed the DeepLabCut output to FMPose3D^40^ to lift the 2D joint trajectories into a frame-by-frame 3D pose sequence from the single video view. We then postprocessed the inferred 3D coordinates in R (v4.5.3; The R Foundation for Statistical Computing, Vienna, Austria) and projected them onto the sagittal plane to generate the stick diagrams shown in Fig. 1A.

### Human body weight suspension system

Low gravity was simulated by means of body weight suspension inside a 3 x 3 m, 17-meter-tall *cavaedium* (light shaft)^12^ at the Locomotion On Other Planets (L.O.O.P.) laboratory, an European Space Agency ground-based facility located in the Human Physiology Building at the University of Milan^41^. The body suspension system consists of 4-meter-long bungee cords (Diagoline S.r.l., Ponderano, BI, Italy) with a stiffness of 92.7 N/m. The bungee cords are linked in series with a short, inextensible 4 mm diameter, 1.2 m long Dyneema SK78 rope (Gottifredi Maffioli S.r.l., Novara, NO, Italy), that slides inside a low-friction upper pulley. One bungee cord is fixed to the wall and the other is connected to a force transducer (TS 300 kg, AEP Transducers, Modena, MO, Italy), which is placed in series with a body-suspension harness. The upper pulley can be raised or lowered by a suspension cable connected to a 2.20 kW motorised winch (Officine Iori S.r.l., Albinea, RE, Italy), which allows unloading the body to the desired gravity level^12^.

### Human simulated hypogravity locomotion and Electromyographic (EMG) recordings

Twelve participants (five females; age: 31.1 ± 5.7 years; height: 1.77 ± 0.07 m; body mass: 69.8 ± 14.1 kg; mean ± SD) were asked to walk, skip, and run at 1.1, 1.4, and 3.1 m/s on a double split-belt instrumented treadmill (FIT, Bertec Corporation, Columbus, OH, United States) at 1.00 g (Earth gravity) and at two simulated gravities: 0.38 g (Martian gravity), and 0.17 g (lunar gravity). Each acquisition lasted 60 s. The order of gaits and gravity levels was randomised. Participants rested for five minutes when the gravity level changed and whenever they requested it. At least 24 hours prior to the experiments, each participant underwent a dedicated familiarisation session in which they became accustomed to the setup, gaits, and gravities. The muscle activity of the following 6 muscles was recorded bilaterally, for a total of 12 muscles, using a 16-channel wireless EMG system (Trigno, Delsys, Natick, MA, United States), with a frequency of 2000 Hz: *vastus medialis* (vm), *semitendinosus* (st), *biceps femoris* (long head, bf), *tibialis anterior* (ta), *gastrocnemius medialis* (gm), and *soleus* (so). Prior to electrode placement each recording area was shaved, abraded, and cleaned with an alcohol wipe/swab^42^. Electrodes were positioned according to SENIAM guidelines^43^. Vertical ground reaction forces were recorded at 2 kHz from the instrumented treadmill, synchronously with EMG. These forces were used to detect the touchdown and take-off of each foot with a vertical threshold of 20 N.

### Muscle synergies extraction

Muscle synergies were extracted through custom scripts (R v4.5.3) using the default parameters of our R package musclesyneRgies^44^ v1.2.5.9009 using the classical Gaussian non-negative matrix factorisation (NMF) algorithm^45^. The raw EMG signals were band-pass filtered within the acquisition device (cut-off frequencies 10 and 500 Hz). Then the signals were high-pass filtered, full-wave rectified and lastly low-pass filtered using a 4^th^ order IIR Butterworth zero-phase filter with cut-off frequencies 50 Hz (high-pass) and 20 Hz (low-pass for creating the linear envelope of the signal). After subtracting the minimum, the amplitude of the EMG recordings was normalised to the maximum activation recorded for each trial for every individual muscle (i.e. every EMG channel was normalised to its maximum in every trial)^18^. Each gait cycle was then time-normalised to 200 points. Synergies were then extracted through NMF. For the analysis, we considered the 12 muscles described above. The m = 12 time-dependent muscle activity vectors were grouped in a matrix **V** with dimensions m × n (m rows and n columns). The dimension n represented the number of normalised time points (i.e. 200*number of gait cycles). The matrix **V** was factorised using NMF so that **V** ≈ **V**_R_ = **WP**. The new matrix **V**_R_, reconstructed multiplying the two matrices **W** and **P**, approximates the original matrix **V**. The matrix **P** contained the time-dependent coefficients of the factorisation (activation patterns) with dimensions r × n, where the number of rows r represents the minimum number of synergies necessary to satisfactorily reconstruct the original set of signals **V**. The matrix **W**, with dimensions m × r, contained the time-invariant muscle weights, which describe the relative contribution of single muscles within a specific synergy (a weight was assigned to each muscle for every synergy). **W** and **P** described the synergies necessary to accomplish the required task (i.e. walking, skipping or running). The update rules for **W** and **P** are:

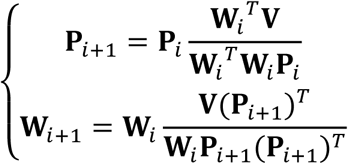

The quality of reconstruction was assessed by measuring the coefficient of determination *R^2^*between the original and the reconstructed data (**V** and **V**_R_, respectively), defined as the ratio between the residuals and total sum of squares^42^:

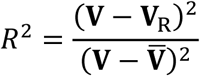

The limit of convergence for each synergy was reached when a change in the calculated *R^2^* was smaller than the 0.01% in the last 20 iterations^42^ meaning that, with that amount of synergies, the signal could not be reconstructed any better. This operation was first completed by setting the number of synergies to 1. Then, it was repeated by increasing the number of synergies each time, until a maximum of 9 synergies. The number 9 was chosen to be lower than the number of muscles, since extracting a number of synergies equal to the number of measured EMG activities would not reduce the dimensionality of the data. Specifically, 9 is the 75% of 12, which is the number of considered muscles^42^. For each synergy, the factorisation was repeated 10 times, each time creating new randomised initial matrices **W** and **P**, in order to avoid local minima^46^. The solution with the highest *R^2^* was then selected for each of the 9 synergies. To choose the minimum number of synergies required to represent the original signals, the curve of *R^2^* values versus synergies was fitted using a simple linear regression model, using all 9 synergies. The mean squared error ^47^ between the curve and the linear interpolation was then calculated. Afterwards, the first point in the *R^2^*-vs.-synergies curve was removed and the error between this new curve and its new linear interpolation was calculated. The operation was repeated until only two points were left on the curve or until the mean squared error fell below 10^−4^. This was done to search for the most linear part of the *R^2^*-versus-synergies curve, assuming that in this section the reconstruction quality could not increase considerably when adding more synergies to the model.

### Muscle synergies analysis

We quantified the temporal structure of muscle synergy activation patterns using full width at half maximum (*FWHM*) and centre of activity (*CoA*) measurements. We calculated the *FWHM* for each gait cycle by subtracting the minimum, normalizing to the peak, and defining it as the number of time-normalised points at which the activation pattern exceeded half of its maximum. The mean *FWHM* across cycles was used for analysis. The *CoA* was computed using circular statistics. For an activation pattern **P** sampled at *p* equally spaced points with phases *θ* ∈ [0, 2π]:

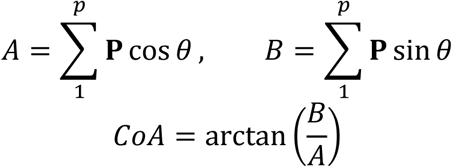

The angle was mapped to the normalised gait cycle (0–100%). To quantify the dispersion of activation timing, we calculated the circular standard deviation:

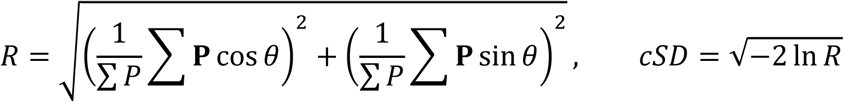

### Cross-reconstruction and permutation analysis

To evaluate how similar muscle synergy structures are across different locomotion conditions, we performed a cross-reconstruction analysis of EMG signals. This analysis used time-independent muscle synergy weights and time-dependent activation patterns. For each subject and gravity level, we reconstructed EMG signals from a reference condition (running) using either the same synergy weights (within-condition reconstruction) or weights from a comparison condition (skipping), combined with the reference condition’s activation patterns. Synergies were treated as four functionally labelled modules functionally classified as explained above. When a labelled synergy was absent in a given trial, its corresponding weight vector was set to zero so that missing synergies would not contribute to the reconstruction. This ensured that all reconstructions were performed in a fixed canonical synergy space, without remapping or interpolation of missing modules. Reconstruction quality between the original data **V** and the reconstructed data **V**_R_ was quantified using the variance accounted for (VAF) defined as

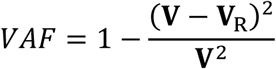

To assess the specificity of synergy ordering, reconstruction was repeated for all 24 possible permutations of the four synergy labels by reordering the weight matrices while keeping the activation patterns unchanged. Permutation selection was performed only on Earth-gravity trials: for each reference comparison, all 24 possible orders of the four comparison-gait weight vectors were tested against the reference-gait activation patterns under Earth gravity. The order that maximised reconstruction quality was then applied, without re-optimisation, to simulated Martian and lunar trials. This prevented the ordering of modules from being defined by locomotor patterns produced in newly experienced hypogravity conditions, which may not have reflected the habitual locomotor repertoire of the participants. It also allowed the hypogravity analyses to test the generalisation of an Earth-derived reference architecture. For each gravity level, the quality of the reconstruction was compared across three types: within-condition reconstruction using the original ordering; cross-reconstruction using the same ordering; and cross-reconstruction using a reordered synergy set.

### Mouse models

We generated *PV^cre^*;*Rx3^flpo^*;*Mapt^dsDTR^*;Ai65D transgenic mice using the following lines. *PV^cre^*: Jackson Laboratories, Bar Harbor, ME, USA; RRID:IMSR_JAX:008069^48^. *Rx3^flpo^*: courtesy of Dr Joriene De Nooij at Columbia University, New York, NY, USA^30^; Ai65D: Jackson Laboratories, Bar Harbor, ME, USA; RRID:IMSR_JAX:021875^31^.

### Mouse body weight suspension system

Hypogravity was simulated with a vertical elastic suspension system that used a solid, round silicone cord (HokoFLEX®, nominal diameter 1.0 mm ± 0.35 mm, 60 ± 5 Shore A, axial stiffness ∼1.7 N/m; HOKOSIL® Elastomertechnik GmbH, Bredenbek, Germany) connected to the animal with a commercially available soft Velcro mouse harness (Butterfly Mouse Harness, Lomir Biomedical Inc., Notre-Dame-de-l’Île-Perrot, QC, Canada). This harness consists of a single piece of back-to-back Velcro with separate neck and thoracic closures, which allow for an adjustable fit while leaving the forelimbs free and the mouth and neck unobstructed. To maintain the line of pull near the anteroposterior location of the mouse centre of mass, an additional Velcro strip was placed between the dorsal harness and the base of the tail. The suspension was designed to provide a constant upward force that corresponds to the desired reduction in effective gravity. To achieve this, the cord rest length was chosen based on the installed vertical span and the mass of the mouse. For the body mass range used in this study and a fixed installed span of 1.50 m, the resulting mean cord extension was approximately 80 to 115 mm under lunar conditions. Kinematic recordings showed a maximum vertical excursion of the centre of mass of about 20 mm. Under these conditions, the expected variation in step-to-step force remained moderate and was primarily determined by the ratio of vertical body oscillation to mean cord extension. Material nonlinearities and hysteresis were neglected at the design stage because force variability was dominated by geometry.

### Open field assay

Open field assay was performed using an ActiMot infrared beam activity monitor (TSE Systems, Berlin, Germany). We analysed a group of 7 adult littermates (three females) two different days, 72 hours apart, first without the harness. Each animal was placed in the arena and allowed to explore freely for 60 minutes. We quantified volitional activity in consecutive 10-minute bins using the manufacturer’s analysis software.

### Estimation of the anteroposterior location of the centre of mass from passive suspension

The experiment was conducted on 4 young adult littermates (2 females; age 152 days; mass 27.9 ± 1.4 g) and 4 older adult littermates (2 females; age 70 days; mass 22.2 ± 0.7 g) mice. After euthanasia, the mice were positioned in a standardised trotting posture, with one diagonal limb pair extended and the other flexed. They were then allowed to undergo *rigor mortis* in this configuration before being stored at -80 °C. Suspension experiments were conducted at 4 °C. Each frozen specimen was suspended from a fixed overhead point via the tail-base strap of the harness in a dorsal position. The specimens were suspended at successive positions spaced 5 mm apart along the full anteroposterior extent of the dorsal strap. At each position, a photograph was taken once the animal reached stable equilibrium. The images were analysed using the ImageJ2 distribution Fiji v2.17.0^49^. A reference segment was drawn from the external auditory meatus to the ventral surface of the tail base. A perpendicular projection from the suspension point onto this segment defined the normalised suspension position. The equilibrium angle relative to horizontal was recorded at each position. The anteroposterior centre-of-mass location was estimated as the zero crossing of a linear regression of body angle on normalised suspension position.

### Mouse treadmill locomotion

Experiments were conducted on 23 adult (13 females; age 73 ± 8 days; mass 23.2 ± 4.7 g) mice. Before treadmill recordings, animals were briefly anesthetised with isoflurane (3% in oxygen, 1 L/min delivered by inhalation). The right hindlimb was shaved and anatomical landmarks were highlighted using a water-based white marker placed over the hip joint and iliac crest to facilitate kinematic tracking. For hypogravity experiments, the body weight support harness was fitted during the same anaesthesia period. Animals were then transferred to the treadmill and allowed to fully recover before data acquisition. Recordings only started once animals resumed spontaneous exploratory behaviour and completed at least one full grooming sequence, indicating recovery from anaesthesia and re-acclimatisation to the experimental environment.

For baseline treadmill locomotion and line characterisation, animals were tested at 0.1, 0.3, 0.5, 0.7, and 0.9 m/s, presented in ascending order. For each speed, we aimed to record at least 100 gait cycles of the right hind leg. This was done in one or any number of trials necessary to reach a total of 100 cycles. After each trial, animals were given a two-minute recovery period before proceeding further. If an animal failed to maintain locomotion at a given speed, the condition was repeated a total of three times. Speeds that could not be successfully maintained after three attempts were classified as not achieved, and no further higher speed conditions were tested for that animal. The animals were tested in the morning.

For simulated hypogravity experiments, the protocol was identical except that recordings were performed only at 0.3 m/s. The body weight support harness was applied during the initial anaesthesia period. The harness was carefully positioned around the thorax and neck while ensuring unrestricted movement of the forelimbs and head. The animals were tested in the morning.

### Motion capture

The kinematics data were recorded through one high-speed camera (boA1936-400cc, Basler AG, Ahrensburg, Germany) operating at 250 Hz. For markerless body part tracking we used DeepLabCut, v2.3.9^24^. We labelled 57 body landmarks and 6 calibration markers on 96 frames taken from 12 videos of 8 different animals, assigning 95% of those images to the training set without cropping. Namely, we labelled six calibration markers from the sagittal view and additionally:

- From the sagittal view: snout, right eye and ear; ankle and metatarsal joints, toe tip of all paws; the right hindlimb iliac crest and hip; scapula, most dorsal point of the trunk, most ventral part on the spine between the previous and the tail base; five equidistant points on the tail from the tail base to the tail tip.
- From the bottom mirror (ventral) view: snout, mouth, ears; the middle and each finger of every paw; five equidistant points on the tail from the tail base to the tail tip.

Iliac crest and hip were highlighted by two white-marker dots as described above. We used a ResNet-50-based neural network^38,39^ with default parameters for 1,100,000 training iterations including two refinements. We validated with one shuffle and found the test error was 4.51 pixels and the train error 2.47 pixels.

### Gait cycle breakdown

Gait events were identified from the filtered kinematic trajectories of the distal limb markers. Specifically, the detection of touchdown and lift-off of each paw relied on the horizontal and vertical motion of the toe tip and metatarsal landmarks from all four limbs. The marker trajectories were low-pass filtered using a fourth-order, zero-phase Butterworth filter with a cut-off frequency of 20 Hz. Trials containing prolonged halts, direction changes, tracking failures, or sustained contact of the tail with the back wall of the treadmill were excluded prior to gait event detection.

For each limb, normalised horizontal toe tip displacement was differentiated to obtain velocity and acceleration profiles. Candidate stride events were first identified from periodic changes in horizontal toe tip motion. Specifically, we detected peaks corresponding to transitions between forward and backward horizontal toe-tip velocity from a more strongly smoothed version of the horizontal displacement signal (8 Hz low-pass filter), with minimum peak spacing adapted to the expected stride frequency. Then, touchdown timing was refined within a narrow temporal window (-5, +20 ms) around each candidate event^18^. The algorithm searched for the earliest peak in vertical acceleration between the metatarsal and toe tip markers. This accounted for variability in foot strike patterns across steps and animals^50^. Additionally, tracking likelihood values from DeepLabCut were used to avoid touchdown assignments during periods of poor marker confidence. Lift-off was identified as the instant of minimum horizontal toe-tip displacement between two consecutive touchdown events, corresponding to the onset of forward paw progression during swing. Touchdown and lift-off timings were calculated independently for all four limbs. To improve cycle consistency and minimise misassignments, gait cycles were aligned relative to the timing of the right hind limb cycle.

### Locomotor-state analysis

To test whether simulated lunar gravity alters the repertoire of accessible mouse locomotor states at baseline, we conducted an unsupervised state-space analysis. Gait cycles were aligned to the right hindlimb cycle. We extracted kinematic descriptors for each cycle to describe temporal gait structure, interlimb coordination, left-right asymmetry, limb endpoint trajectories, airtime, support geometry, axial body motion, trunk and hindlimb joint kinematics, and cycle-to-cycle variability. We normalised length-dependent variables by femur-plus-tibia length and base-of-support area by the square of this length scale. We retained metadata including mouse identity, gravity condition, trial identity, and cycle index for downstream analysis, but did not use them as state-defining variables.

We calculated unsigned left-right asymmetry as |L − R|/(|L| + |R|) and signed asymmetry as (L − R)/(|L| + |R|). Hind-fore relative contrasts were calculated as (hind − fore)/(|hind| + |fore|). Interlimb phase-coupling variables were computed as the circular distance between touchdown phases, expressed relative to the right hindlimb cycle. Cycle-to-cycle dynamic descriptors were calculated as the relative change between consecutive right hindlimb cycles. Descriptors with excessive missing data were removed, and cycles with excessive missing data were excluded. The remaining missing descriptor values were imputed using the median value of the corresponding descriptor across the retained dataset. The descriptors were then centred and scaled, and principal component analysis was used to obtain a low-dimensional representation of the cycle-level kinematic state space. Locomotor states were inferred across Earth- and simulated lunar-gravity cycles using Gaussian mixture modelling on the retained principal component scores so that the same state labels could be used in both conditions. State identity was characterised *post hoc* using descriptor fingerprints and representative cycles. State accessibility was quantified from the inferred state sequence by calculating state occupancy, entry probability, dwell time, and self- and row-normalised transition probabilities between states. Hidden Markov modelling of the same principal component scores was used for temporal sensitivity analysis.

### Diphtheria toxin administration

Each adult *PV^cre^*;*Rx3^flpo^*;*Mapt^dsDTR^*;Ai65D mouse received an intraperitoneal injection of diphtheria toxin (D0654 lyophilised; Sigma-Aldrich, St. Louis, MO, USA), diluted in ultrapure water at a concentration of 100 μg/kg^26,27^. The final solution contained 10 μg/ml of toxin. For example, a 32-g mouse would receive 320 μl of the solution.

### Immunohistochemistry

For immunohistochemistry analysis, each *PV^cre^*;*Rx3^flpo^*;*Mapt^dsDTR^*;Ai65D mouse was perfused after deep anaesthesia was achieved using a diluted cocktail of xylazine and ketamine in 1X phosphate buffered saline (PBS). For every 10 g of mouse body mass, the cocktail was prepared with 20 µl of 10% ketamine (Ketabel®, 100 mg/ml), 16 µl of 2% xylazine (Rompun®, 20 mg/ml), and 64 µl of 1X PBS. After thoracotomy, the mice were perfused with 20 ml of 1X PBS followed by 10 ml of 4% paraformaldehyde (PFA) solution through the left cardiac ventricle. The tissue of interest was dissected and post-fixed in 4% PFA at 4 °C overnight. Then, it was cryoprotected by immersion in a 30% sucrose-1X PBS solution for at least 24 hours or until it sank. After cryoprotection, the tissue was embedded in an optimal cutting temperature (OCT) mounting medium, flash frozen on dry ice, and stored at -80 °C. The tissue was sectioned using a cryostat (Leica CM3050 S, Leica Biosystems AG, Muttenz, Switzerland) and placed on microscope slides to dry for at least one hour. The muscle, brainstem, and brain samples were sectioned at 50 µm, the dorsal root ganglia at 25 µm, and the spinal cord at 20 µm. For all types of tissue, we initially washed the slides three times in 1X PBS for 10 minutes each to remove OCT. Then, the slides were incubated overnight at 4 °C in a primary antibody solution diluted in 1X PBS + 0.1% Triton X-100 (PBT). The next day, the slides were washed three more times in 1X PBT for five minutes each. Then, the slides were incubated for one hour at room temperature with a secondary antibody solution in PBT. After three more 5-minute washes in 1X PBS, the slides were mounted with Mowiol mounting medium and coverslipped. Primary antibodies: guinea pig anti-parvalbumin (1:3000, Swant, GP72), guinea pig anti-VGLUT1 (1:2000, Sigma-Aldrich, AB5905), and rabbit anti-DsRed (1:1000, Takara, 632496). Secondary antibodies: Alexa Fluor 488 donkey anti-guinea pig IgG (H+L) (1:500, Jackson ImmunoResearch, 706-545-148) and Cy3 donkey anti-rabbit IgG (H+L) (1:500, Jackson ImmunoResearch, 711-165-152). NeuroTrace 435 (1:500, Invitrogen, N21479) was used as a fluorescent Nissl stain.

### Statistics

The statistical analyses were performed in R v4.5.3 using linear mixed-effects models, unless otherwise stated. Participants were included as random intercepts to account for repeated measurements within individuals. For outcomes measured repeatedly across synergy, identity, or reconstructed objects, the corresponding term was included as an additional random intercept when required by the analysis. Fixed effects were selected according to the design of each analysis. For outcomes measured across gait and gravity, such as interlimb step-time asymmetry, the number of extracted synergies, and electromyography (EMG) reconstruction quality, models included gait, gravity, and their interaction as fixed effects. The interlimb asymmetry analysis was restricted to the gait-speed combinations shown in Fig. 1E: walking at 1.1 m/s, skipping at 1.4 m/s, and running at 3.1 m/s. For the short-term maximum Lyapunov exponent of activation-pattern dynamics, models were fitted separately at each gravity level, with gait as the fixed effect. For the full width at half maximum (FWHM) of activation patterns, models were fitted separately for each gait-speed condition. These models included gravity, synergy identity, and their interaction as fixed effects and participant and synergy object as random intercepts.

Fixed effects were evaluated using Type III tests. The Kenward-Roger degrees-of-freedom approximation was used for analyses implemented with Kenward-Roger mixed-model inference. Type III Wald tests were used for models fitted with Wald inference. Post hoc comparisons were performed on estimated marginal means. Tukey adjustment was used for all-pair comparisons within a single contrast family. Holm adjustment was used for targeted families of contrasts, such as gait comparisons within each gravity level or gravity comparisons within each gait level. Compact letter display^51^ was used as graphical summaries of adjusted *post hoc* comparisons. For each comparison family, the estimated marginal means were ordered from lowest to highest before letters were assigned, with “a” denoting the lowest estimated value and subsequent letters denoting progressively higher values. Conditions that shared at least one letter were not statistically distinguishable at the 0.05 level of significance, whereas conditions that had no letters in common differed significantly. Thus, the letters indicate the ordinal direction of the effect under this ordering convention but not its magnitude.

For analyses of left-right limb asymmetry in mouse, asymmetry values were bounded between 0 and 1. These values were squeezed away from the exact boundaries and logit-transformed before statistical modelling. For intact mice, asymmetry was analysed with gravity condition, limb pair, and their interaction as fixed effects. For proprioceptor ablation experiments, day after diphtheria toxin injection was included as an additional fixed effect, together with its interactions with gravity condition and limb pair. Pairwise contrasts between gravity conditions were computed within limb pair and day using estimated marginal means and Holm correction for multiple testing. Compact letter displays were used to summarise the adjusted post hoc comparisons at α = 0.05.

Principal component analyses of mouse kinematics were performed using separate linear mixed-effects models for scores on PC1 and PC2. The fixed effect was the experimental factor represented in the corresponding figure: either the gravity condition or the day after diphtheria toxin injection. Mouse identity was included as a random intercept. Pairwise contrasts among days were adjusted using the Holm method. When testing PC1 and PC2 together, the resulting families of p-values were adjusted using Holm correction across principal components. For the effect-size analysis of lunar-gravity-induced hindlimb asymmetry, the condition contrasts were standardised using the residual standard deviation from the corresponding mixed-effects model. They were then reported as Cohen’s d on the transformed scale. Confidence intervals were obtained from the estimated marginal contrasts.

For the harness-control experiment, distance travelled and average moving speed were analysed with harness condition as the fixed effect and mouse identity as the random intercept. Histological quantification of proprioceptor ablation was analysed with the number of days since diphtheria toxin injection as a fixed effect. When cell-level measurements were used, mouse identity was included as a random intercept to account for the nesting of cells within animals. Model assumptions were evaluated by inspecting residual distributions and fitted versus residual plots.

## Acknowledgments

The authors are grateful to Liana Kosizki for technical support, to Feride Salman, Sonya Yung, Marta Sanchez Acedo, Cristian Bonomo, and Paula Lindenberg for supporting mouse tissue preparation and imaging; to the Animal Phenotyping platforms at the MDC for supporting mouse behavioural setups, and to Turgay Akay, Graziana Gatto, Quinn Silverman, and members of the Zampieri laboratory for providing comments on the manuscript.

## Author contributions

Conceptualisation: AS, AEM, GP, NZ Methodology: AS, AEM, GP, NZ Investigation: AS, FL, VN, AM, NM, GP Visualisation: AS Funding acquisition: AEM, GP, NZ Project administration: AEM, GP, NZ Supervision: AS, AEM, GP, NZ Writing – original draft: AS, NZ Writing – review & editing: all authors

## Data, code, and materials availability

All primary data have been deposited in Zenodo at https://doi.org/10.5281/zenodo.20394408. All unique reagents and materials generated in this study are available from the authors upon reasonable request.

## Figure legends

**Fig. S1.**
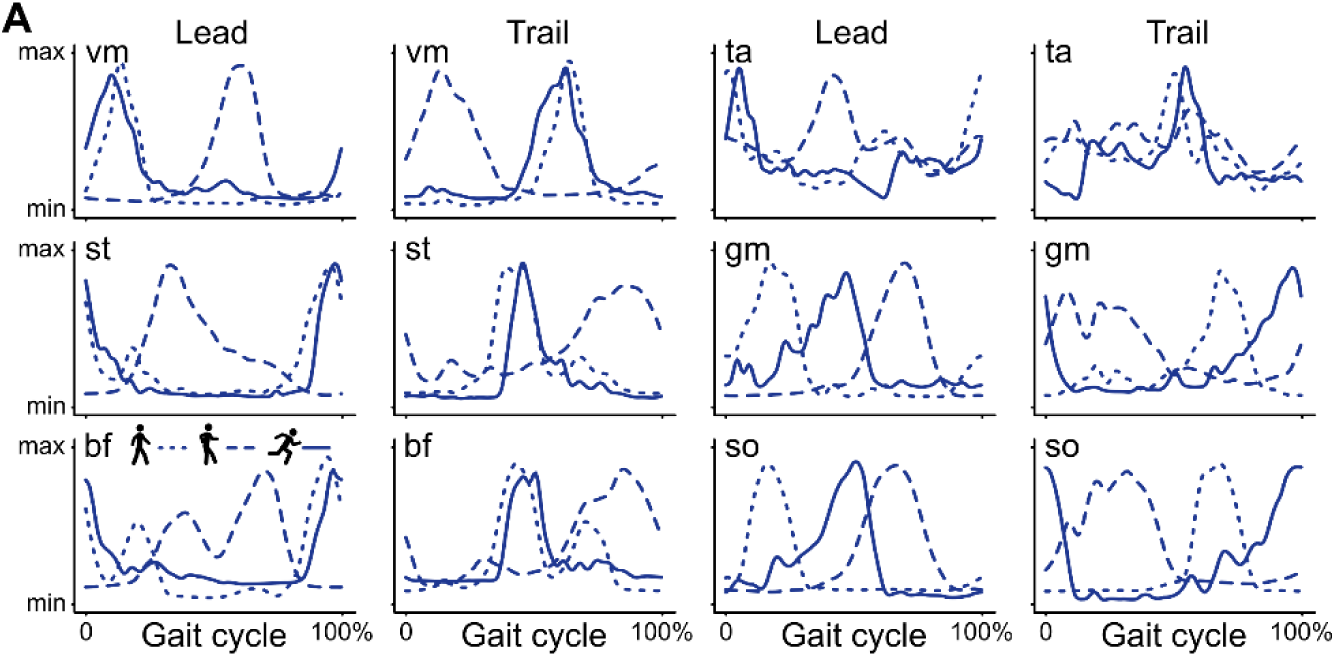
Lower-limb muscle activity across gait modes under Earth gravity. (**A**) Average electromyographic activity profiles for individual muscles during walking at 1.1 m/s, skipping at 1.4 m/s, and running at 3.1 m/s under Earth gravity (*N* = 12 participants). Signals are shown separately for the leading and trailing legs, time-normalized to the gait cycle, and amplitude-normalized between minimum and maximum activation. Line type indicates gait mode, as shown by the icons. Abbreviations: vm = *vastus medialis*; st = *semitendinosus*; bf = *biceps femoris*; ta = *tibialis anterior*; gm = *gastrocnemius medialis*; so = *soleus*.

**Fig. S2.**
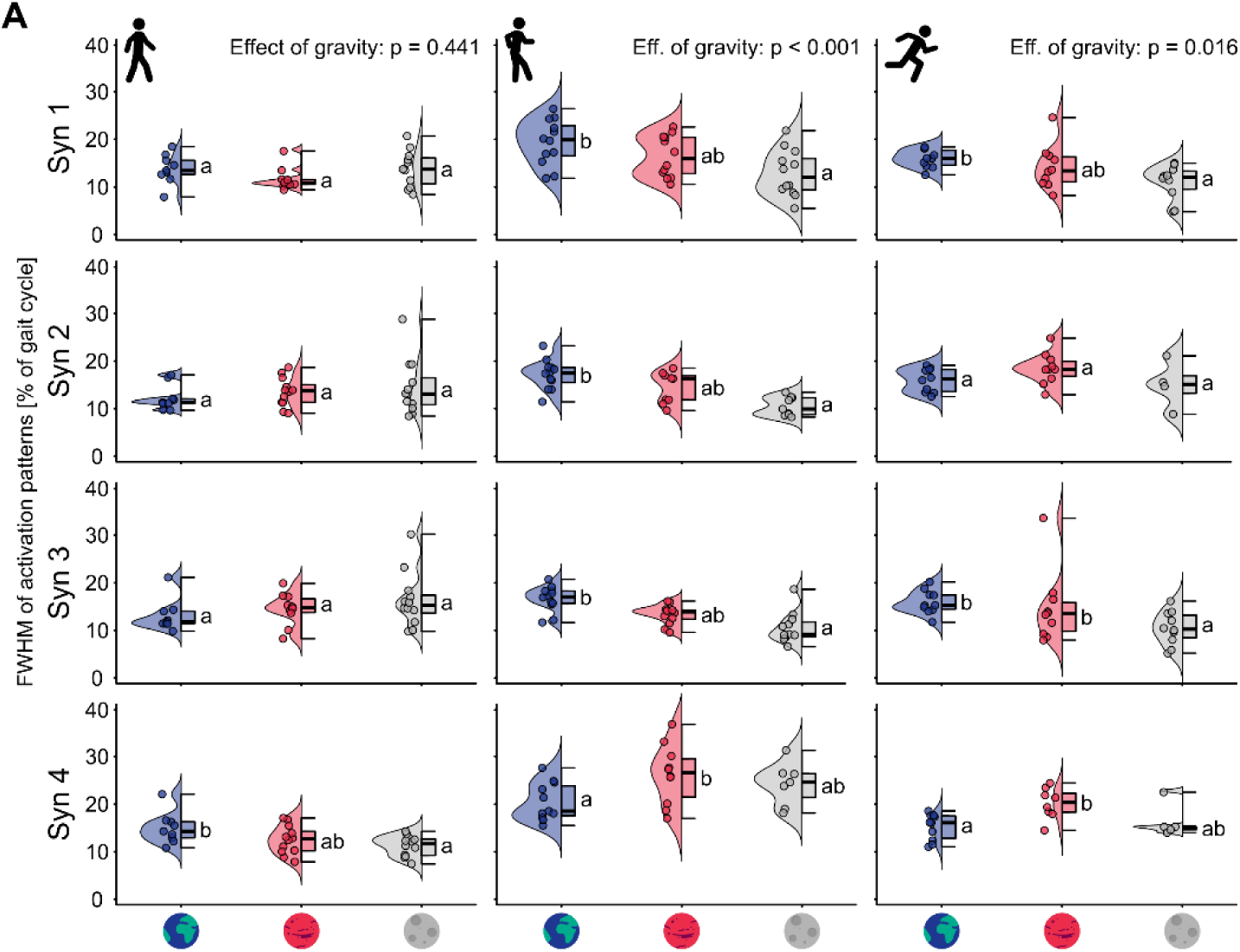
Temporal width of muscle-synergy activation patterns across gaits and gravity levels. (**A**) Full width at half maximum (FWHM) of synergy activation patterns for walking at 1.1 m/s, skipping at 1.4 m/s, and running at 3.1 m/s under Earth, Martian, and lunar gravity. Rows show individual synergies (Syn1 to Syn4), and columns show gait modes. Within each panel, distributions across gravity conditions are shown as violin plots, with overlaid boxplots and individual trial estimates. FWHM values are expressed as percentage of the gait cycle. Different letters indicate significant *post-hoc* differences (*N* = 12 participants).

**Fig. S3.**
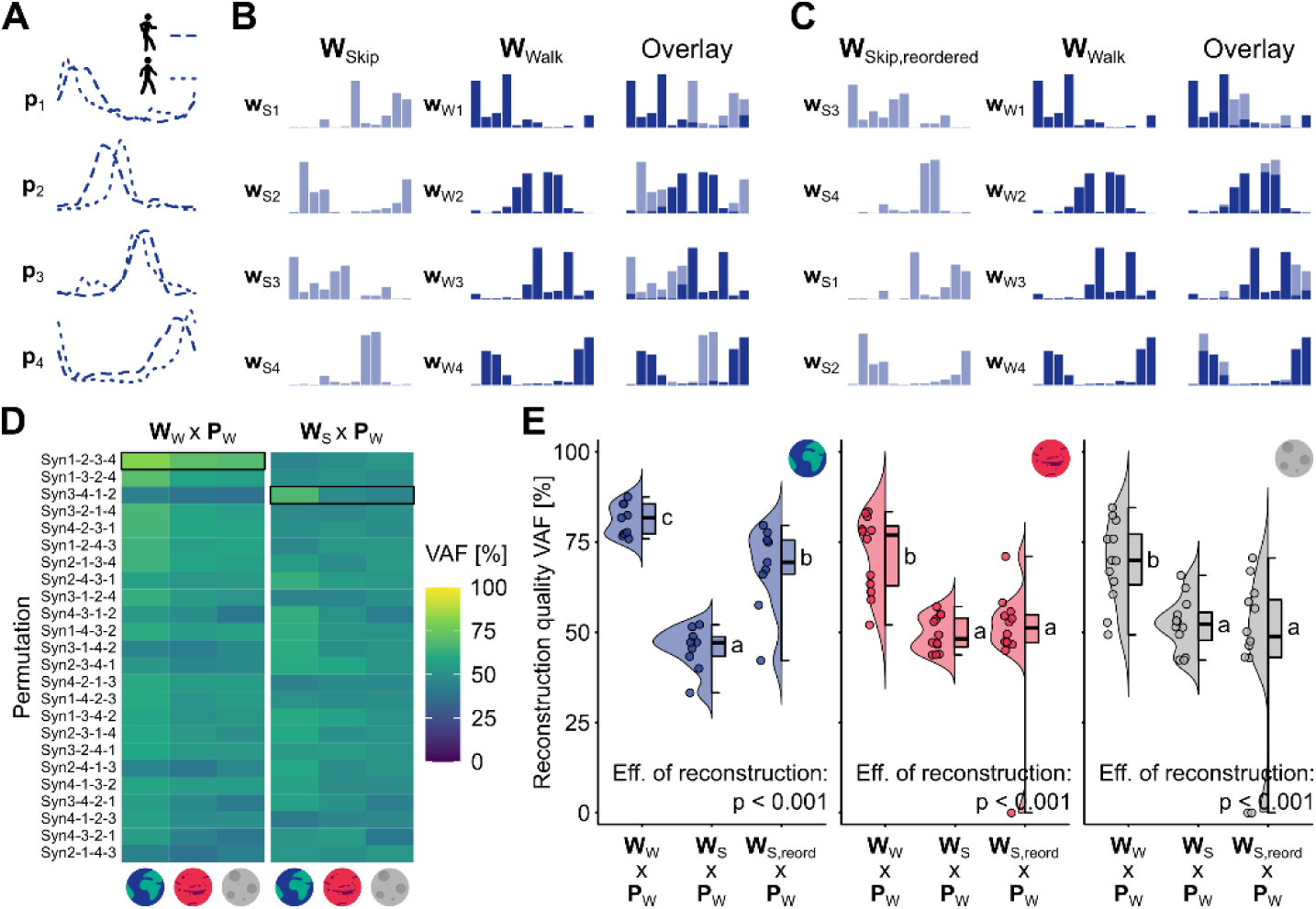
Skipping does not share a walking-like modular architecture. (**A**) Average synergy activation patterns **P** for walking and skipping under Earth gravity. (**B**) Synergy muscle weights **W** for walking and skipping under Earth gravity. (**C**) Reordered skipping muscle weights overlaid on walking weights. (**D**) Cross-reconstruction analysis of running EMG using walking activation patterns combined with walking weights or with skipping weights. Reconstruction quality was quantified for all possible weight-order permutations. (**E**) Reconstruction quality of walking across gravity conditions using i) walking weights and activations, ii) skipping weights and walking activations, or iii) reordered skipping weights and walking activations. Different letters indicate significant *post-hoc* differences (*N* = 12 participants).

**Fig. S4.**
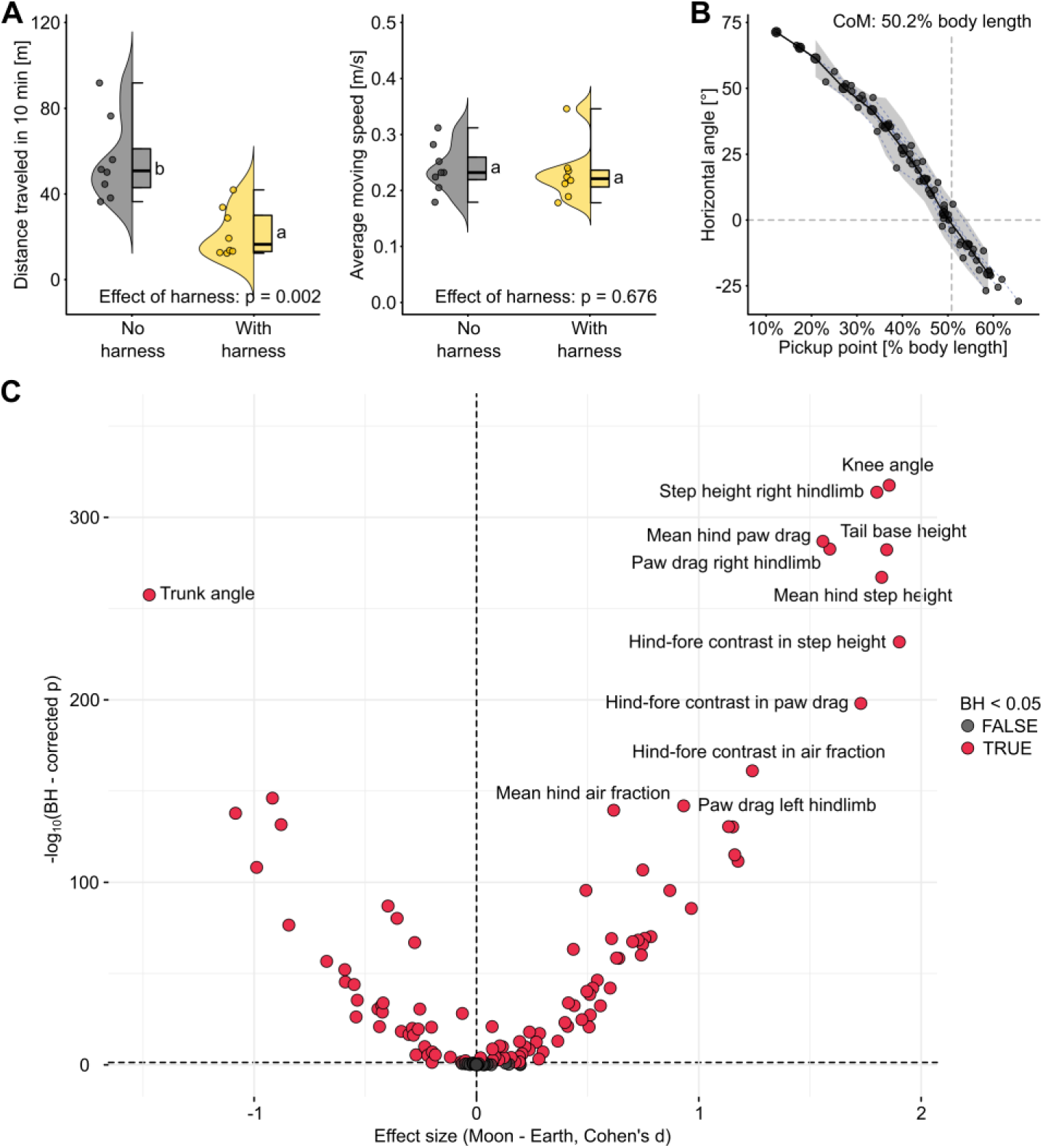
Effects of the body-weight support harness on mouse locomotion. (**A**) Distance traveled and average moving speed of mice in open field test with or without the body-weight support harness (*N* = 8 mice). Different letters denote significant *post-hoc* differences. (**B**) Estimation of the anteroposterior position center of mass of the mouse from passive suspension tests (*N* = 8 mice; *n* = 4 young adults; *n* = 4 older adults). Frozen mice were positioned in a standardized trotting posture and suspended from successive pickup points along the dorsal harness strap. The horizontal body angle was plotted as a function of pickup-point distance from the tail base, expressed as a percentage of body length. The thin lines and small dots show the data for individual mice, the thick line and big dots show the mean across mice, and the shaded ribbon indicates ±1 standard deviation. The horizontal dashed line marks zero body angle, and the vertical dotted line indicates the interpolated zero crossing used to estimate the anteroposterior center-of-mass position. Different letters indicate significant *post-hoc* differences. (**C**) Volcano plot summarizing the difference between simulated lunar gravity and Earth gravity across all extracted kinematic descriptors. Each point represents one descriptor. The x-axis shows the standardized effect size, expressed as Cohen’s *d* for Moon minus Earth, so positive values indicate larger descriptor values under simulated lunar gravity and negative values indicate larger values under Earth gravity. The y-axis shows statistical evidence for the Moon versus Earth difference, expressed as −log10 of the Benjamini-Hochberg-corrected *p-*value. Red points indicate descriptors that remained significant after false-discovery-rate correction, whereas grey points indicate non-significant descriptors. Labelled points highlight a few selected descriptors with large effect sizes and/or strong statistical evidence.

**Fig. S5.**
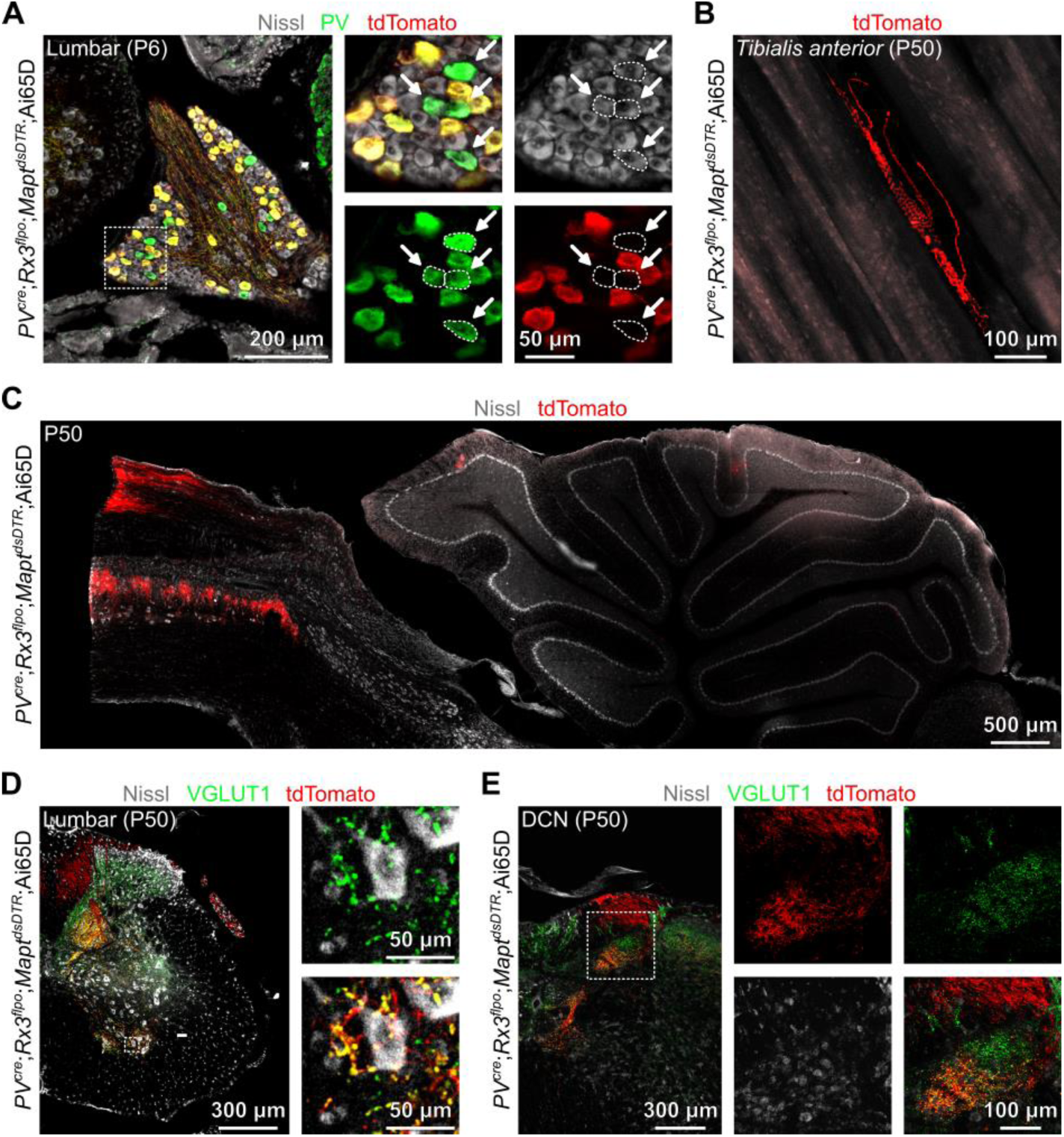
Anatomical validation of proprioceptor targeting. (**A**) A lumbar dorsal root ganglion at P6 showing Nissl staining, parvalbumin (PV) immunolabeling, and tdTomato expression. The insets show the overlap between PV and tdTomato in sensory neurons. As expected, not all PV-positive neurons express tdTomato **(*26*, *50*)**. (**B**) tdTomato-labeled muscle spindle in the tibialis anterior muscle at P50. (**C**) Sagittal view of tdTomato-labeled proprioceptive afferents in the cervical spinal cord and hindbrain at P50. (**D**) Coronal section of lumbar spinal cord at P50 shows tdTomato-labeled proprioceptive afferents and co-localization with VGLUT1-positive presynaptic puncta on motor neurons. (**E**) Coronal view of the hindbrain at P50 show tdTomato-labeled proprioceptive afferents projections to the dorsal column nuclei (DCN).

**Fig. S6.**
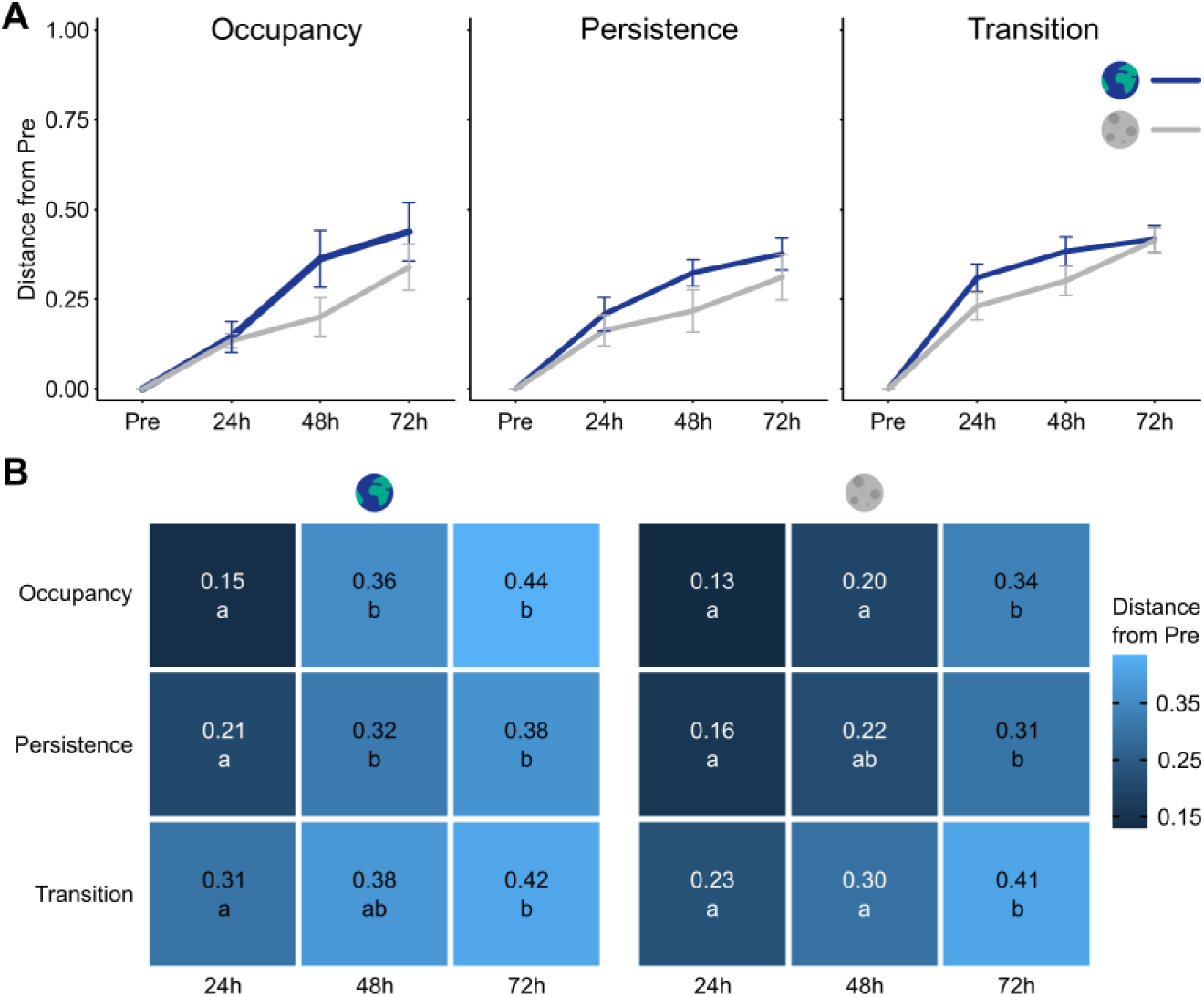
Locomotor-state landscape divergence from pre-exposure baseline. (**A**) Bray-Curtis distance of the locomotor-state landscape from the pre-exposure baseline, computed separately for occupancy, persistence, and transition vectors. “Pre” (i.e. the baseline before DT injection) was used as the reference and therefore has distance 0 by definition. Lines show group means and error bars indicate mean ± SE for Earth gravity controls and simulated lunar gravity. (**B**) Compact heatmap summary of landscape divergence from “Pre”. Cell color indicates the estimated Bray-Curtis distance from “Pre”, and numbers show the corresponding estimated distance. Letters indicate compact-letter-display groupings among 24 h, 48 h, and 72 h within each gravity condition and landscape metric; time points sharing a letter are not significantly different from each other, whereas time points with non-overlapping letters differ in their distance from “Pre”. The heatmap is intended to summarize the magnitude and temporal progression of whole-landscape divergence, rather than state-by-state changes in individual occupancy, persistence, or transition probabilities.

**Table S1.**
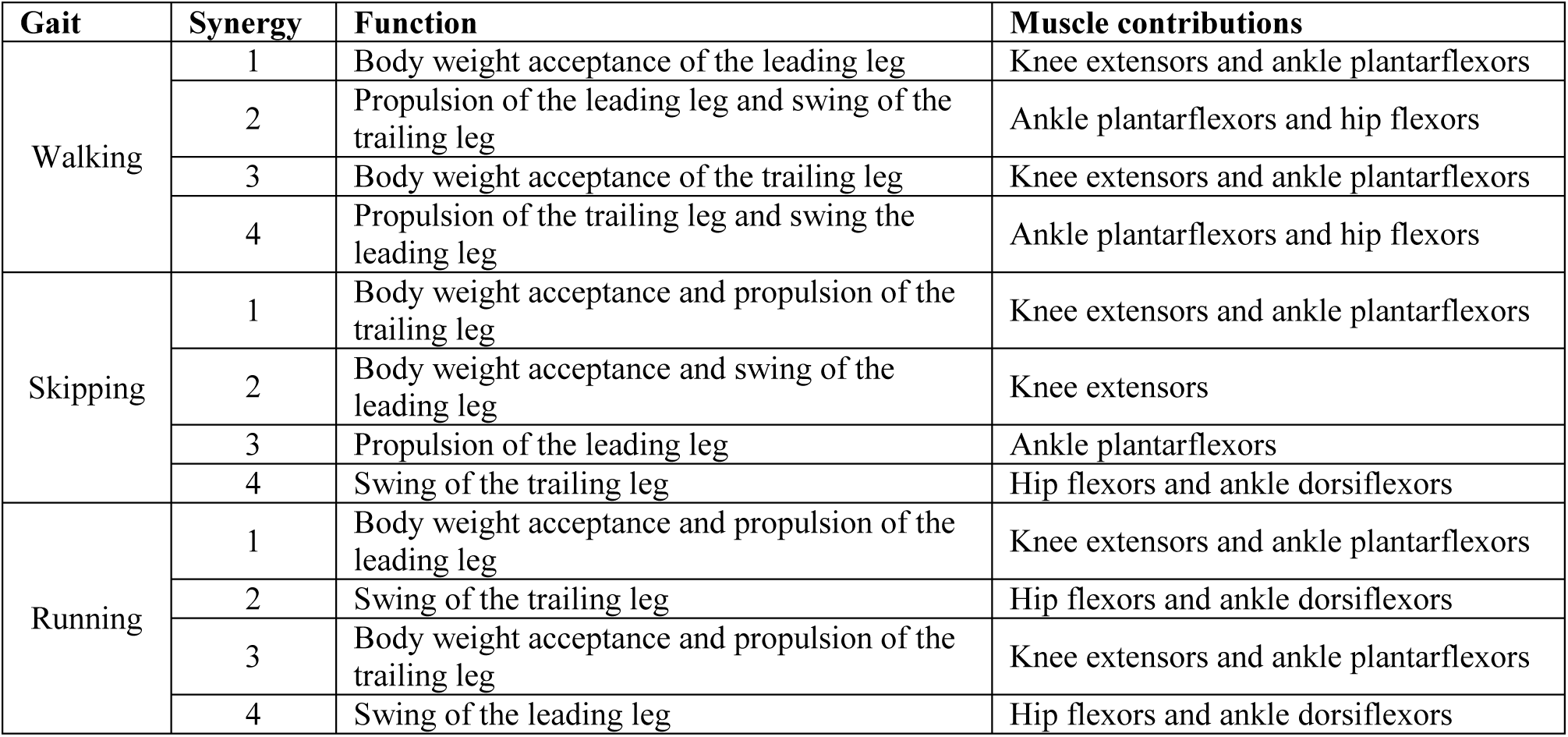
Function of muscle synergies during walking, skipping, and running. Muscle synergies extracted from EMG activity in the lower limbs during walking, skipping and running were determined based on the timing of their activation patterns within the gait cycle and the relative contribution of individual muscles to the corresponding muscle weights.

**Table S2.** Kinematic descriptors included in the mouse locomotor-state analysis. Kinematic descriptors (134 in total) were extracted from markerless motion tracking during treadmill locomotion under both Earth’s and simulated lunar gravity. The table lists descriptor families, units before centering and scaling, and operational definitions. Unless otherwise stated, per-limb variables were calculated for the left forelimb, right forelimb, left hindlimb, and right hindlimb. Bilateral forelimb and hindlimb means, left-right asymmetries, signed asymmetries, hind-to-fore relative contrasts, and selected cycle-to-cycle changes were derived from these primary variables. Metadata such as mouse and trial identities, gravity condition, and cycle index were retained for downstream analysis but were not used as state-defining variables.

| Category | Parameters | Unit | Definition |
| --- | --- | --- | --- |
| Temporal gait structure | Cadence | cycles/min | 60 divided by stride duration between consecutive touchdowns of the same limb. |
|  | Stance duration | s | Time from touchdown to subsequent lift-off of the same paw. |
|  | Swing duration | s | Time from lift-off to subsequent touchdown of the same paw. |
|  | Duty factor | dimensionless | Stance duration divided by stride duration. |
|  | Forelimb and hindlimb bilateral means | source-variable unit | Mean of the left and right limb values within the forelimb or hindlimb pair. |
| | Hind-fore temporal contrasts | dimensionless | Relative contrast between hindlimb and forelimb bilateral means, calculated as $(\text{hind} - \text{fore}) / (\text{abs}(\text{hind}) + \text{abs}(\text{fore}))$. |
| Interlimb coordination and left-right asymmetry | Touchdown phase | gait-cycle fraction | Touchdown timing relative to right-hindlimb touchdown, divided by right-hindlimb stride duration and wrapped to the interval [0, 1). |
|  | Hindlimb and forelimb left-right phase distances | gait-cycle fraction | Circular distance between left and right hindlimb phases, or between left and right forelimb phases. |
|  | Diagonal phase distances | gait-cycle fraction | Circular distance between left forelimb and right hindlimb phases, and between right forelimb and left hindlimb phases. |
|  | Homolateral phase distances | gait-cycle fraction | Circular distance between forelimb and hindlimb phases on the same body side. |
| | Phase-coupling asymmetry | dimensionless | Unsigned asymmetry between the two diagonal or the two homolateral phase distances, calculated as $\text{abs}(A - B) / (\text{abs}(A) + \text{abs}(B))$. |
| | Unsigned left-right asymmetry | dimensionless | Absolute left-right difference normalized by signal magnitude, calculated as $\text{abs}(L - R) / (\text{abs}(L) + \text{abs}(R))$. |
| | Signed left-right asymmetry | dimensionless | Signed left-right difference normalized by signal magnitude, calculated as $(L - R) / (\text{abs}(L) + \text{abs}(R))$. |
| Limb endpoint trajectories and airtime | Stride length | dimensionless | Anteroposterior range of the toe-tip trajectory over one gait cycle, divided by femur plus tibia length. |
|  | Step height | dimensionless | Vertical range of the toe-tip trajectory over one gait cycle, divided by femur plus tibia length. |
|  | Paw drag | a.u. | Area under the vertical toe-tip trajectory during swing, after subtracting the local minimum paw height and normalizing vertical displacement by femur plus tibia length. |
|  | Air fraction | dimensionless | Fraction of frames within the gait cycle in which the toe-tip vertical position was above a limb-specific trial threshold. |
|  | Forelimb and hindlimb bilateral means | source-variable unit | Mean of the left and right limb endpoint or airtime values within the forelimb or hindlimb pair. |
| | Hind-fore endpoint contrasts | dimensionless | Relative contrast between hindlimb and forelimb bilateral means for stride length, step height, paw drag, and air fraction, calculated as $(\text{hind} - \text{fore}) / (\text{abs}(\text{hind}) + \text{abs}(\text{fore}))$. |
| Support geometry | Base of support | dimensionless area | Mean convex-hull area defined by stance paw landmarks during the right-hindlimb gait cycle, divided by (femur + tibia length)<sup>2</sup>. |
| Axial-body kinematics | Tail-base height | dimensionless | Minimum tail-base vertical position relative to the lowest right-hindlimb toe-tip position within the same cycle, divided by femur plus tibia length. |
|  | Tail-base vertical excursion | dimensionless | Vertical range of the tail-base trajectory during the right-hindlimb gait cycle, divided by femur plus tibia length. |
|  | Tail-base horizontal excursion | dimensionless | Anteroposterior range of the tail-base trajectory during the right-hindlimb gait cycle, divided by femur plus tibia length. |
|  | Tail-base path length | dimensionless | Two-dimensional path length of the tail-base trajectory during the right-hindlimb gait cycle, divided by femur plus tibia length. |
|  | Sagittal trunk angle mean and range of motion | deg | Mean and range of motion of the sagittal trunk angle during the right-hindlimb gait cycle. |
| Hindlimb joint kinematics | Hip angle mean and range of motion | deg | Mean and range of motion of the hip angle during the right-hindlimb gait cycle. |
|  | Knee angle mean and range of motion | deg | Mean and range of motion of the knee angle during the right-hindlimb gait cycle. |
|  | Ankle angle mean and range of motion | deg | Mean and range of motion of the ankle angle during the right-hindlimb gait cycle. |
|  | Metatarsophalangeal angle mean and range of motion | deg | Mean and range of motion of the metatarsophalangeal angle during the right-hindlimb gait cycle. |
| Cycle-to-cycle variability and dynamics | Ataxia coefficient | dimensionless | Trial-level stride-length range for a limb divided by that limb's current-cycle stride length. |
|  | Bilateral mean ataxia coefficient | dimensionless | Mean of the left and right ataxia coefficients within the forelimb or hindlimb pair. |
|  | Left-right ataxia asymmetry | dimensionless | Unsigned and signed left-right asymmetry of the ataxia coefficient. |
|  | Hind-fore ataxia contrast | dimensionless | Relative contrast between hindlimb and forelimb bilateral mean ataxia coefficients, calculated as (hind - fore) / (abs(hind) + abs(fore)). |
|  | Relative cycle-to-cycle changes | dimensionless | Relative change between consecutive cycles, calculated as (current - previous) / (abs(current) + abs(previous)), for selected descriptors. |
|  | Circular phase change | gait-cycle fraction | Absolute circular change in hindlimb left-right phase distance between consecutive cycles. |

**Movie S1. Eugene Cernan’s locomotion on the lunar surface.**

Original NASA Apollo 17 video footage showing Eugene Cernan moving on the lunar surface. The sequence illustrates naturally occurring locomotor strategies in lunar gravity, including transitions between running, skipping, and hopping-like patterns. The MPEG video was obtained from the Apollo 17 Lunar Surface Journal.

**Movie S2. Human walking, skipping and running in simulated lunar gravity.**

Representative participant walking, skipping, and running on a treadmill under simulated lunar gravity. Walking is shown on the left at 1.1 m/s, skipping in the middle at 1.4 m/s, and running on the right at 3.1 m/s. Footprint schematics indicate the corresponding footfall sequence and follow the convention used in Fig. 1C.

**Movie S3. Mouse treadmill locomotion under Earth and simulated lunar gravity.**

Representative mouse locomoting on a treadmill at 0.3 m/s under Earth gravity, left, and simulated lunar gravity, right. Videos are shown at 20% real speed.

**Movie S4. Mouse treadmill locomotion before and 48 hours after proprioceptor ablation under Earth gravity.**

Representative mouse treadmill locomotion under Earth gravity at 0.3 m/s before ablation, left, and 48 hours after proprioceptor ablation, right. Videos are shown at 20% real speed.

**Movie S5. Mouse treadmill locomotion 72 hours after proprioceptor ablation.**

Representative mouse locomoting on a treadmill at 0.3 m/s 72 hours after proprioceptor ablation under Earth gravity, left, and simulated lunar gravity, right. Videos are shown at 20% real speed.

**Movie S6. Mouse treadmill locomotion before and 48 hours after proprioceptor ablation under simulated lunar gravity.**

Representative mouse treadmill locomotion under simulated lunar gravity at 0.3 m/s before ablation, left, and 48 hours after proprioceptor ablation, right. Videos are shown at 20% real speed.

## References

1. Frigon, A., Akay, T. & Prilutsky, B. I. Control of Mammalian Locomotion by Somatosensory Feedback. Comprehensive Physiology 12, 2877–2947 (2021).

2. Goulding, M., Bollu, T. & Büschges, A. Sensory Feedback and the Dynamic Control of Movement. Annual Review of Neuroscience 48, 405–423 (2025).

3. Kiehn, O. Decoding the organization of spinal circuits that control locomotion. Nature Reviews Neuroscience 17, 224–238 (2016).

4. Lacquaniti, F. et al. Human Locomotion in Hypogravity: From Basic Research to Clinical Applications. Front. Physiol. 8, 893 (2017).

5. De Martino, E., Green, D. A., Ciampi De Andrade, D., Weber, T. & Herssens, N. Human movement in simulated hypogravity—Bridging the gap between space research and terrestrial rehabilitation. Front. Neurol. 14, 1062349 (2023).

6. Akay, T., Tourtellotte, W. G., Arber, S. & Jessell, T. M. Degradation of mouse locomotor pattern in the absence of proprioceptive sensory feedback. Proceedings of the National Academy of Sciences of the United States of America 111, 16877–82 (2014).

7. Minetti, A. E. The biomechanics of skipping gaits: a third locomotion paradigm? Proc. R. Soc. Lond. B 265, 1227–1233 (1998).

8. Jones, E. M. & Glover, K. Apollo 17 Lunar Surface Journal: Station 5. (2017).

9. Jones, E. M. & Glover, K. Apollo 14 Lunar Surface Journal: Traverse to Station F. (2017).

10. Mission operations branch & Flight crew support division. Apollo 11 Technical Crew Debriefing Vol. I. https://www.nasa.gov/history/alsj/a11/a11tecdbrf.html (1969).

11. Ivanenko, Y. P., Cappellini, G., Poppele, R. E. & Lacquaniti, F. Spatiotemporal organization of á-motoneuron activity in the human spinal cord during different gaits and gait transitions. European Journal of Neuroscience 27, 3351–3368 (2008).

12. Pavei, G., Biancardi, C. M. & Minetti, A. E. Skipping vs. running as the bipedal gait of choice in hypogravity. Journal of Applied Physiology 119, 93–100 (2015).

13. National Aeronautics and Space Administration. NASA’s Lunar Exploration Program Overview. (2020).

14. Bizzi, E., Cheung, V. C.-K., D’Avella, A., Saltiel, P. & Tresch, M. C. Combining modules for movement. Brain Research Reviews 57, 125–33 (2008).

15. Oliveira, A. S. C., Gizzi, L., Kersting, U. G. & Farina, D. Modular organization of balance control following perturbations during walking. Journal of Neurophysiology 108, 1895–906 (2012).

16. Ting, L. H. et al. Neuromechanical Principles Underlying Movement Modularity and Their Implications for Rehabilitation. Neuron 86, 38–54 (2015).

17. Martino, G. et al. Neuromuscular adjustments of gait associated with unstable conditions. Journal of Neurophysiology 114, 2867–2882 (2015).

18. Santuz, A., Ekizos, A., Eckardt, N., Kibele, A. & Arampatzis, A. Challenging human locomotion: stability and modular organisation in unsteady conditions. Scientific Reports 8, 2740 (2018).

19. Brüll, L. et al. Spatiotemporal modulation of a common set of muscle synergies during unpredictable and predictable gait perturbations in older adults. Journal of Experimental Biology 227, jeb.247271 (2024).

20. Santuz, A. & Akay, T. Muscle spindles and their role in maintaining robust locomotion. The Journal of Physiology 601, 275–285 (2023).

21. Duysens, J., Clarac, F. & Cruse, H. Load-regulating mechanisms in gait and posture: Comparative aspects. Physiological Reviews 80, 83–133 (2000).

22. Santuz, A. et al. Neuromotor Dynamics of Human Locomotion in Challenging Settings. iScience 23, 100796 (2020).

23. Lyapunov, A. M. The general problem of the stability of motion. International Journal of Control 55, 531–534 (1992).

24. Mathis, A. et al. DeepLabCut: markerless pose estimation of user-defined body parts with deep learning. Nature Neuroscience 21, 1281–1289 (2018).

25. Bellardita, C. & Kiehn, O. Phenotypic Characterization of Speed-Associated Gait Changes in Mice Reveals Modular Organization of Locomotor Networks. Current Biology 25, 1426–1436 (2015).

26. Takeoka, A. & Arber, S. Functional local proprioceptive feedback circuits initiate and maintain locomotor recovery after spinal cord injury. Cell Reports 27, 71–85.e3 (2019).

27. Santuz, A., Laflamme, O. D. & Akay, T. The brain integrates proprioceptive information to ensure robust locomotion. The Journal of Physiology 600, 5267–5294 (2022).

28. de Nooij, J. C., Doobar, S. & Jessell, T. M. Etv1 Inactivation Reveals Proprioceptor Subclasses that Reflect the Level of NT3 Expression in Muscle Targets. Neuron 77, 1055–1068 (2013).

29. Wu, D. et al. A Role for Sensory end Organ-Derived Signals in Regulating Muscle Spindle Proprioceptor Phenotype. J. Neurosci. 39, 4252–4267 (2019).

30. Oliver, K. M. et al. Molecular correlates of muscle spindle and Golgi tendon organ afferents. Nature Communications 12, (2021).

31. Britz, O. et al. A genetically defined asymmetry underlies the inhibitory control of flexor–extensor locomotor movements. eLife 4, e04718 (2015).

32. Hayes, G. & Alexander, R. McN. The hopping gaits of crows (Corvidae) and other bipeds. Journal of Zoology 200, 205–213 (1983).

33. Moore, T. Y., Cooper, K. L., Biewener, A. A. & Vasudevan, R. Unpredictability of escape trajectory explains predator evasion ability and microhabitat preference of desert rodents. Nat Commun 8, 440 (2017).

34. Verstappen, M., Aerts, P. & Damme, R. V. Terrestrial Locomotion in the Black-Billed Magpie: Kinematic Analysis of Walking, Running and Out-of-Phase Hopping. Journal of Experimental Biology 203, 2159–2170 (2000).

35. Wunderlich, R. E. & Schaum, J. C. Kinematics of bipedalism in *Propithecus verreauxi*. Journal of Zoology 272, 165–175 (2007).

36. Proske, U. & Gandevia, S. C. The Proprioceptive Senses: Their Roles in Signaling Body Shape, Body Position and Movement, and Muscle Force. Physiological Reviews 92, 1651–1697 (2012).

37. Grillner, S. & El Manira, A. Current Principles of Motor Control, with Special Reference to Vertebrate Locomotion. Physiological Reviews 100, 271–320 (2020).

38. He, K., Zhang, X., Ren, S. & Sun, J. Deep residual learning for image recognition. Proceedings of the IEEE Computer Society Conference on Computer Vision and Pattern Recognition 2016-Decem, 770–778 (2016).

39. Insafutdinov, E., Pishchulin, L., Andres, B., Andriluka, M. & Schiele, B. DeeperCut: A Deeper, Stronger, and Faster Multi-person Pose Estimation Model. in European Conference on Computer Vision (eds Leibe, B., Matas, J., Sebe, N. & Welling, M.) vol. 9910 34–50 (Springer International Publishing, Cham, 2016).

40. Wang, T., Yu, X. & Mathis, M. W. FMPose3D: monocular 3D pose estimation via flow matching. Preprint at 10.48550/ARXIV.2602.05755 (2026).

41. Herssens, N. et al. Movement in low gravity environments (MoLo) programme–The MoLo-L.O.O.P. study protocol. PLoS ONE 17, e0278051 (2022).

42. Santuz, A., Ekizos, A., Janshen, L., Baltzopoulos, V. & Arampatzis, A. On the Methodological Implications of Extracting Muscle Synergies from Human Locomotion. International Journal of Neural Systems 27, 1750007 (2017).

43. Stegeman, D. F. & Hermens, H. J. Standards for surface electromyography: the European project ‘Surface EMG for non-invasive assessment of muscles (SENIAM)’. 108–112 (1999).

44. Santuz, A. musclesyneRgies: factorization of electromyographic data in R with sensible defaults. Journal of Open Source Software 7, 4439 (2022).

45. Lee, D. D. & Seung, H. S. Learning the parts of objects by non-negative matrix factorization. Nature 401, 788–91 (1999).

46. D’Avella, A. & Bizzi, E. Shared and specific muscle synergies in natural motor behaviors. Proceedings of the National Academy of Sciences of the United States of America 102, 3076–81 (2005).

47. Cheung, V. C.-K., D’Avella, A., Tresch, M. C. & Bizzi, E. Central and sensory contributions to the activation and organization of muscle synergies during natural motor behaviors. The Journal of Neuroscience 25, 6419–34 (2005).

48. Hippenmeyer, S. et al. A developmental switch in the response of DRG neurons to ETS transcription factor signaling. PLoS Biology 3, 0878–0890 (2005).

49. Schindelin, J., et al. Fiji: an open-source platform for biological-image analysis. Nat Methods 9, 676–682 (2012).

50. Santuz, A. & Akay, T. Fractal analysis of muscle activity patterns during locomotion: pitfalls and how to avoid them. Journal of Neurophysiology 124, 1083–1091 (2020).

51. Piepho, H.-P. An Algorithm for a Letter-Based Representation of All-Pairwise Comparisons. Journal of Computational and Graphical sStatistics 13, 456–466 (2004).

